# Modeling the emergent metabolic potential of soil microbiomes in Atacama landscapes

**DOI:** 10.1101/2024.12.23.630026

**Authors:** Constanza M. Andreani-Gerard, Natalia E. Jiménez, Ricardo Palma, Coralie Muller, Pauline Hamon-Giraud, Yann Le Cunff, Verónica Cambiazo, Mauricio González, Anne Siegel, Clémence Frioux, Alejandro Maass

## Abstract

**Background:** Soil microbiomes harbor complex communities and exhibit important ecological roles resulting from biochemical transformations and microbial interactions. Difficulties in characterizing the mechanisms and consequences of such interactions together with the multidimensionality of niches hinder our understanding of these ecosystems. The Atacama Desert is an extreme environment that includes unique combinations of stressful abiotic factors affecting microbial life. In particular, the Talabre Lejía transect has been proposed as a unique natural laboratory for understanding adaptation mechanisms.

**Results:** We propose a systems biology-based computational framework for the reconstruction and simulation of community-wide and genome-resolved metabolic models, in order to provide an overview of the metabolic potential as a proxy of how microbial communities are prepared to respond to the environment. Through a multifaceted approach that includes taxonomic and functional profiling of microbiomes, simulation of the metabolic potential, and multivariate analyses, we were able to identify key species and functions from six contrasting soil samples across the Talabre Lejía transect. We highlight the functional redundancy of whole metagenomes, which act as a gene reservoir from which site-specific functions emerge at the species level. We also link the physicochemistry from the puna and the lagoon samples to specific metabolic machineries that could be associated with their adaptation to the unique environmental conditions found there. We further provide an abstraction of community composition and structure for each site that allows to describe them as sensitive or resilient to environmental shifts through putative cooperation events.

**Conclusion:** Our results show that the study of community-wide and genome-resolved metabolic potential, together with targeted modeling, may help to elucidate the role of producible metabolites in the adaptation of microbial communities. Our framework was designed to handle non-model microorganisms, making it suitable for any (meta)genomic dataset that includes nucleotide sequence data and high-quality environmental metadata for different samples.

## Background

Soil bacterial communities are particularly heterogeneous and complex (Sokol et al., 2022). They demonstrate specific functional responses to their environment, and can modify their surroundings by passively releasing or actively secreting metabolites (Pande and Kost, 2017). Bacterial communities also exhibit self-organizing properties by sharing labor costs to adapt to specific environmental constraints such as nutrient availability, for example through synergistic interactions (Anantharaman et al., 2016; Thommes et al., 2019; Mataigne et al., 2021).

From a ecological perspective, it is generally assumed that beneficial interactions within bacterial communities are based on species engaging in synergistic behavior, including metabolic exchange between species (Ziesack et al., 2019; Louis et al., 2014). Several studies suggest that metabolic exchanges allow for the re-utilization of metabolites released into the environment, benefiting not only the producer but also neighboring microorganisms. These molecules have been referred to as “public goods” (Boon et al., 2014) and could explain the evolution of functional capabilities in members of these communities. This behavior, often referred to as “metabolic handoffs” (Hug and Co, 2018), and the intrinsic metabolic dependencies in crossfeeding determine not only microbial community composition, but also its stability, as the properties of microorganisms compensating for others’ gene loss affect the ability of the whole community to overcome perturbations (Mataigne et al., 2021; Morris et al., 2012; Shade et al., 2012). In line with this, the ecological concept of keystone species, a term originally defined in the context of food web complexity and community stability (Paine, 1969), corresponds to members whose removal can cause a dramatic change in the structure and function of the microbiome (Wang et al., 2024).

Functions harboured by, possibly low-abundant, keystone species are context-dependent and predicted to be critical for the ecosystem, as they catalyze load points in community-wide metabolic networks (Muller et al., 2018). Thus, metabolism, the first response of the microbiome to environmental perturbations, is a critical level of analysis to better understand the assembly of the community (Rothman et al., 2023; van der Knaap and Verrijzer, 2016; Kochanowski et al., 2017; Wang and Lei, 2018). However, it has been hypothesized that the high diversity of microbial communities, especially those of soils, favors functional redundancy where different taxa can perform the same set of metabolic processes and, therefore, can easily replace each other (Allison and Martiny, 2008). Although efforts have been made to unravel the relationships between nutrient cycling processes and the architecture of soil microbiomes at the taxonomic and functional levels (Zhou et al., 2022), providing mechanistic insights into the relationship between nutrient availability and microbiome composition remains a puzzling challenge. We still do not fully understand how community-level functional properties in the microbiome emerge from the assimilation and transformation of environmental nutrients. This requires deciphering everything from how the metabolic machinery of the community is activated in response to the nutritional properties of the culture medium, to the metabolites secreted by the community, and how this metabolite production is characteristic of the physicochemical properties of the medium inhabited by the community. (Silverstein et al., 2024).

Metabolic modeling has been shown to be a successful approach to account for cross-feeding and other potential microbial interactions in whole communities when applied to metagenomic information, looking at the metabolism of the community as a whole or analyzing its individuals through metagenome-assembled genomes (MAGs) (Lambert et al., 2024; Budinich et al., 2017; Taş et al., 2021; Xun et al., 2021; Régimbeau et al., 2022). However, the latter is difficult to achieve, as the extensive microbial diversity prevents the reconstruction of genomes for the majority of low abundance populations, especially in complex ecosystems such as soil (Ejaz et al., 2024). Our main hypothesis in this context, is that building metabolic models at both the metagenome and MAG scales can provide relevant hypotheses about the system. Our main focus is to assess whether the metabolic capacities of different species could be linked to different subsets of available niches, and whether the larger metabolic repertoire of the multispecies community allows it to occupy a wider range of niches. In this scenery, metagenome-scale models can be representative of the consequences of metabolic handoffs, as they combine the metabolic capabilities of the entire community (Saleem et al., 2019), while MAG-level models are likely to highlight the metabolism of dominant members of the microbiome within microenvironments of soil (Fierer, 2017).

In this article, we aimed to provide answers to the questions raised above by exploring genomescale metabolism modeling of an entire microbiome and inferring the ability of its networks to produce metabolites of interest (Belcour et al., 2020). To achieve our goal, we take advantage of metagenomic data and measurements of physicochemical conditions collected at six sites along the altitudinal gradient of the Talabre-Lejía transect (TLT), located on the eastern margin of the Salar de Atacama to the Lejía lagoon. This transect captures three vegetation belts: prepuna (2,400 to 3,300 meters above sea level; masl); puna (3,300 to 4,000 masl); and steppe under the influence of a variety of abiotic conditions, and exhibits a level of diversity and heterogeneity that is representative of complex soil microbiomes (Mandakovic et al., 2023; Eshel et al., 2021). Microorganisms in this environment have to withstand extreme conditions of UV radiation, salinity, high diurnal temperature variation and extremely low availability of nutrients and water (Díaz et al., 2016; Mandakovic et al., 2018; Andreani-Gerard et al., 2024). We build on the knowledge gained about this specific ecosystem and analyze it through the lens of systems biology. We develop and apply a metabolic modeling strategy to MAGs and metagenomes assembled from the TLT, in order to unravel the theoretical capacity of its microbes to synthesize metabolites under different environmental conditions. Our results suggest that the relationships between nutrients, metabolic potential and taxa in the transect are site-dependent, despite a common reservoir of functions at the metagenome level. We demonstrate the utility of a strategy that combines community-wide and genome-resolved metabolic models to suggest important pathways and key players in complex microbiomes.

## Methods

### Geographical locations, sample collection and metagenomic sequencing

Soil sampling and extraction analytical protocols, DNA extraction and sequencing from six sites along the Talabre-Lejía Transect (TLT; 23.4°S, 67.8°W) have recently been described (Andreani-Gerard et al., 2024)). Briefly, bulk soil samples (100 g) were collected in triplicate at a depth of 10 cm from the ground spanning an altitudinal gradient with different vegetation cover: pre-puna (S1, 2,400 to 3,300 m.a.s.l.), puna (S2, 3,200 to 4,000 m.a.s.l.), and steppe (S3 to S6, 4,000 to 4,500 m.a.s.l.); and stored on dry ice until arrival at the laboratory for metagenomic sequencing. DNA extraction was performed using the NucleoSpin Food kit (Macherey-Nagel). DNA from triplicates was pooled to obtain one representative DNA sample per site. Sequencing was performed by MR DNA (www.mrdnalab.com, Shallowater, TX, USA) on a Miseq platform (Illumina, San Diego, CA) with an overlapping 2×150 bp configuration. Soil physicochemical measurements are presented in Table S1.

Nearly 453 million paired-end reads of 150 nucleotides in length were generated across the six sampled sites. In total, 325,603,002 of the nearly 430 million high quality reads were mapped to the assembled metagenomic contigs (76%), averaging more than 50 million reads per sample (Table S2).

### Metagenome and MAG assembly

Metagenomic reads from the six sequenced samples were trimmed using the BBTools protocols (Bushnell et al., 2017) to remove Illumina adapters and low quality bases. High quality reads (>90%) were then used to build the metagenomic assemblies for each sample using MEGAHIT v1.2.9 (Li et al., 2015) with the kmer preset “meta-large” recommended for soils. Statistics of the assemblies are available in Table S2. A multi-process pipeline was prepared to extract sitespecific metagenome-assembled genomes (MAGs). A triple binning and consensus approach was used, with Maxbin2 v2.2.7 (Wu et al., 2015), Metabat2 v2.2.15 (Kang et al., 2019) and Concoct v1.1.0 (Alneberg et al., 2014). All binners were set to a minimum contig size of 2000 bp and the coverage mapping recommended by Metabat2. To combine and refine all outputs, the Metawrap pipeline (Uritskiy et al., 2018) was used as the consensus method with parameter set to their default recommended values. Resulting bins were quality-assessed using CheckM v1.2.2 (Parks et al., 2015) and filtered to 136 high quality MAGs (completeness >70% and contamination <10%). Taxonomic classification was performed using the GTDB release 214 with its toolkit v2.3.0 (Chaumeil et al., 2019). Dereplication of MAGs classified to the same species was performed following the dRep scoring metric (Olm et al., 2017) with default values for parameters related to completeness, contamination, strain heterogeneity, N50 and size, and with F set to 0. The genomic dataset was reduced to 120 high-quality, unique MAGs. Relative abundance of MAGs was calculated individually using the metagenomic reads belonging to their site of origin, using the CoverM v0.7.0 protocol (https://github.com/wwood/CoverM).

### Taxonomic and functional profiling

Taxonomic classification of the reads mapped to the metagenomic contigs was performed using the mOTUs microbial profiler v3.1 with default settings (Ruscheweyh et al., 2022).

Structural gene annotation in the assembled metagenomes and MAGs was performed using Prodigal v2.6.3 (Hyatt et al., 2010). Functional annotation of these genes was performed using eggNOG-mapper v2.1.6 (Huerta-Cepas et al., 2019) based on eggNOG orthology data release 5.0 (Huerta-Cepas et al., 2018), with sequence searches performed using DIAMOND v2.1.8 (Buchfink et al., 2014). Functional categories from COG (Galperin et al., 2021), PFAM (Finn et al., 2006), CAZYme (Drula et al., 2022) and KEGG (Kanehisa et al., 2024) were assigned for analysis. Genbank files containing annotations and sequences were generated using inhouse scripts based on the SeqIO Biopython library release 1.83 (Cock et al., 2009) for use in downstream metabolic reconstruction.

### Metabolic network reconstruction and modeling

The input to the metabolic network reconstruction was the collection of genbank files described above. Reconstruction was performed using the GeMeNet pipeline (https://gitlab.inria. fr/slimmest/gemenet) based on Pathway-tools v.25.5 (Karp et al., 2022), mpwt v0.7 (Belcour et al., 2020) and Padmet v5.0.1 (Aite et al., 2018).

Simulations of producible metabolites, i.e., the metabolic modeling, were performed with Menetools v3.3.0 (Aite et al., 2018) for metagenomes and Metage2Metabo v1.5.2 (Belcour et al., 2020) for MAGs, using the subcommands `scope` and `metacom` respectively. Both tools require a list of available nutrient compounds, referred to as *seeds*, which initialize the inference of other reachable, i.e., producible, metabolites in the network. This step is referred to as *network expansion* and, in the genome-resolved approach, it takes into account the metabolic complementarity of MAGs, therefore suggesting the producibility of new metabolites resulting from putative cross-feeding interactions. We describe as *MetaG-GEM* a metabolic model obtained from a metagenome-scale metabolic network, and as *MAG-GEM*, a metabolic model resulting from a collection of MAG-scale metabolic networks. Thus, 12 systems were developed: 6 MetaG-GEMs and 6 MAG-GEMs, one of each per site.

Five conditions were designed to simulate the systems, each described as a list of nutrient compounds. The first condition is *basal medium*, comprising inorganic carbon sources (CO_2_ and HCO_3_), water, oxygen, inorganic phosphorus, nitrate, nitrite, ammonium, sulfate, sulfite, hydrogen peroxide, arsenate, molybdate, metal ions (Fe^2+^, Mg^2+^, Ni^2+^, Co^2+^ and Cu^2+^) and other coenzymes and cofactors. This basal medium is included in all four additional conditions. The *simple sugars* condition contains carbohydrates that enter glycolysis before 2-phosphoglycerate, as defined in “Group A” by Wang *et al*. (Wang et al., 2019) (e.g., glucose, maltose, galactose, arabinose, sorbitol and glycerol). The *complex sugars* condition includes soil organic matter such as the molecules obtained in an untargeted metabolomics effort by Swenson et al. (2015) (e.g., trehalose, sucrose, rhamnose, mannitol, xylitol, linoleic acid, spermidine, coumarate and chorismate). The *all amino acids* condition includes the twenty genomically-encoded amino acids. Finally, the *non-sulfured amino acids* condition is the latter set excluding the sulfur-containing amino acids (cysteine and methionine). The conditions are summarized in Table 1 and their detailed composition is available in Table S8.

**Table 1:** Summary of the five conditions constructed for simulations in terms of compounds available for initializing the network expansion. Conditions are detailed further in Table S8.

|  | basal<br>medium | simple<br>sugars | complex<br>sugars | non-sulf.<br>amino acids | all<br>amino acids |
| --- | --- | --- | --- | --- | --- |
| **bioavailable nutrients** |  |  |  |  |  |
| organic carbon | 0 | glucose, maltose,<br>galactose, +[4] | rhamnose, xylitol,<br>spermidine, +[18] | ser, pro, val, thr, ile, leu, gln,<br>lys, his, phe, arg, tyr, gly, ala,<br>asp, asn, glt, trp |  |
| organic nitrogen | 0 | 0 | 0 |  |  |
| organic sulfur | 0 | 0 | 0 | 0 | cys, met |
| subtotal | 0 | 7 | 21 | 18 | 20 |
| **basal medium** |  |  |  |  |  |
| inorganic carbon |  |  | HCO<sub>3</sub>, CO<sub>2</sub> |  |  |
| inorganic nitrogen |  |  | N<sub>2</sub>, ammonium, nitrate, nitrite |  |  |
| inorganic sulfur |  |  | HS, SO<sub>3</sub>, S<sub>2</sub>O<sub>3</sub>, sulfate |  |  |
| other inorg. chem. |  |  | H<sub>2</sub>O, O<sub>2</sub>, H<sub>2</sub>, H<sup>+</sup>, H<sub>2</sub>O<sub>2</sub>, Pi, Cl<sup>-</sup>, +[2] |  |  |
| metal ions |  |  | Mg, Fe, Ni, Co, Cu, MoO<sub>4</sub> |  |  |
| coenzymes |  |  | NAD, FAD, CoA, thiamine, +[14] |  |  |
| subtotal | 43 | 43 | 43 | 43 | 43 |
| **total** | **43** | **50** | **64** | **61** | **63** |

In total, 60 simulations were performed, each of the 12 systems with the 5 conditions, generating predictions of producible metabolites for each. Each simulation therefore resulted in a binary vector describing the producibility (1) or non-producibility (0) of each metabolite in each system and condition. In particular, metabolites predicted in MAG-GEMs simulations that were never predicted in MetaG-GEMS simulations were removed to eliminate the effect of likely gap-filled reactions during network reconstruction step (n=82). The Metage2Metabo and Menetool log files containing the output lists of producible compounds were converted with an in-house script into binary matrices indicating the presence or absence of metabolites in each simulation. Identical presence/absence vectors of metabolites were collapsed into unique occurrence groups of metabolites across simulations (Table S6).

### Statistical analyses

For taxonomic profiling, relative abundances obtained from mOTUs (see *Taxonomic and functional profiling*) were used. OTU data were rarefied to the least sequenced sample using the `rarefy_even_depth` function from the phyloseq package v1.46.0 (McMurdie and Holmes, 2013) for calculating diversity within microbiomes only. Alpha diversity indices were calculated using the `diversity` and `fisher.alpha` functions from the vegan package v2.6 (Oksanen et al., 2020) in R. For functional profiling, the number of annotated genes from annotation reports (see *Taxonomic and functional profiling*) was normalized by the total number of COG, PFAM, KEGG pathway and CAZyme entries detected per sample (Table S5). To determine the taxonomic ranks and functional categories that contributed most to the differences in abundance between the six soil microbiomes, a similarity percentage analysis (SIMPER) was performed using the Bray-Curtis dissimilarity matrix calculated on the Hellinger-transformed data in PAST v4.03 (Øyvind Hammer et al., 2001).

For multivariate analyses, principal Coordinate Analyses (PCoA) were performed with `stats::cmdscale` using the Bray-Curtis dissimilarity matrix calculated on the Hellinger-transformed taxonomic and functional abundances and using the Jaccard index on the binary matrices obtained in previous section. Unsupervised clustering was drawn with `stats_ellipse` (type = “t”, level = 0.95) using the metabolite matrix. Hierarchical clustering of the simplified metabolic data (i.e., metabolite groups, see *Metabolic network reconstruction and modeling*) was performed using the Jaccard index and the ward.D2 linking method, and visualized using the ComplexHeatmap v2.18.0 package (Gu, 2022) in R. For environmental metadata, a principal component analysis (PCA) of soil physicochemical measurements (pH, electrical conductivity (EC), percent organic matter (OM), and fourteen others in mg/kg: N, NH_4_, NO_3_, P, K, Mg, Ca, Cu, Fe, Zn, Mn, B, S and Na) was performed with `stats::prcomp` and plotted with `fviz_pca_biplot` from the factoextra package v1.0.7 in R (Team, 2019).

Lastly, using the matrix of unique occurrence groups of metabolites obtained across simulations, an elastic net regressions were applied with an alpha value of 0.85 using the glmnet package v4.18 (Tay et al., 2023) in R. To elucidate which metabolic traits better predict environmental metadata, the physicochemical measurements obtained *in situ* were individually targeted as explained variables (see Figure 2D for illustration). Unique occurrence groups of metabolites with nonzero coefficients and, thus, fitted as relevant for the regression model, were defined as ‘key’ if absolute values of coefficients were greater than 0.3. Hierarchical clustering of this processed data was performed with the Bray-Curtis dissimilarity distance and the ward.D2 linking method, and plotted with the ComplexHeatmap package in R. For a visual summary of our bioinformatic pipeline, refer to Fig. 2. Lists of metabolites of interest were extracted and used for visualization using Ontosunburst v1.0.0 (https://github.com/AuReMe/Ontosunburst).

All statistical analyses were performed using R v4.3.2. Plots were generated using ggplot v3.5.0 (Wickham, 2016) unless otherwise indicated.

### Selection of minimal communities for the synthesis of targeted metabolites

Metage2Metabo (Belcour et al., 2020) permits selecting minimal communities, i.e., a set of metabolic networks of minimal cardinality that sustain the reachability from nutrients of a set of targeted compounds. We used this approach to identify MAGs that could be involved in the production of environmentally-driven groups of metabolites detected with the elastic net regression model (see *Statistical analyses*). Metage2Metabo’s `mincom` command was executed for each of the six sites providing a single list of all such metabolites as targets. Runs were performed with *basal medium* because selected metabolic groups were unambiguous across seeds, i.e., they displayed the same producibility profiles across simulations regardless of the seed used.

Metage2Metabo provides by default a unique minimal community, together with information relative to the key species, essential and alternative symbionts. The former contains all metabolic networks appearing in at least one of the minimal-size communities. Essential symbionts appear in every minimal community, highlighting possible producibility bottlenecks in the microbiome, whereas alternative symbionts appear in one but not all minimal communities, suggesting functional redundancy for the targeted functions within the original microbial composition. Enumeration of all minimal-size communities ensuring the reachability of metabolic targets was additionally performed, in order to retrieve information related to the co-presence of taxa in the predicted minimal communities. This association of symbionts was visualized with power graph compression as provided in Metage2Metabo. Power graphs were drawn in SVG format with the command `m2m_analysis::workflow`, and styled with Inkscape v1.3.2 to reflect the taxonomic classification and abundance of MAGs.

## Results

### Environmentally unique sampling sites at the Talabre-Lejía transect reflect distinct taxonomic and functional profiles of metagenomes

The Talabre-Lejía transect (TLT, ∼23.5°S) is an altitudinal gradient located from the eastern margin of the Atacama Salar up to the Lejía lagoon that has been extensively studied for capturing different ecosystems and uncover key processes associated with adaptation to the Atacama Desert, the most arid nonpolar environment on Earth (Arroyo et al., 1988; Latorre et al., 2002; Eshel et al., 2021; Mandakovic et al., 2023). The aridity of the TLT is maximal in its lowest altitudes, whereas the highest ones may receive rain in the first three months of the year. Soil samples of our study span the prepuna, puna, and steppe (Fig. 1A).

**Figure 1:**
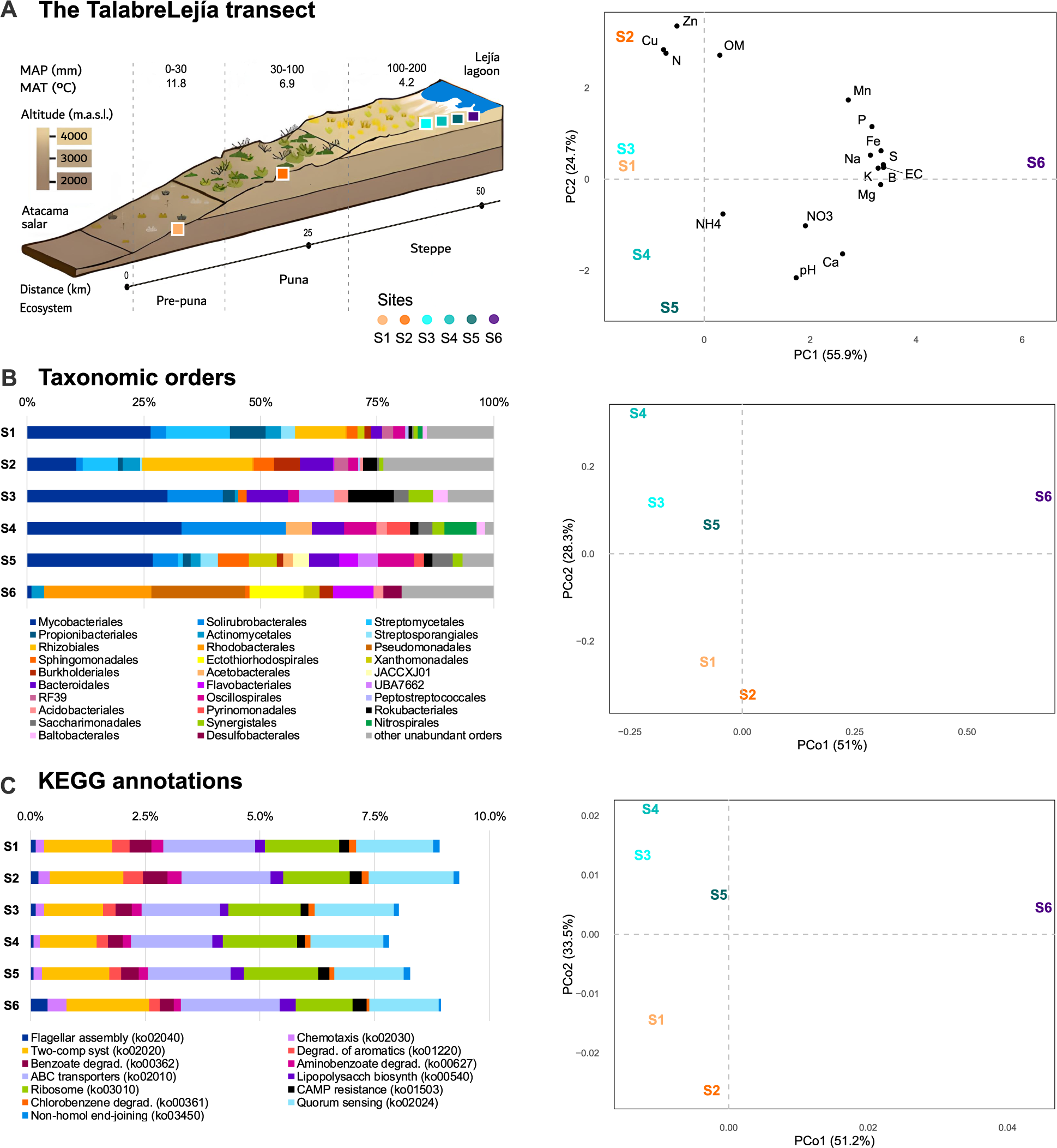
Environmental heterogeneity across the Talabre-Lejía transect (TLT). **A.** Geographic illustration of the TLT adapted from Mandakovic et al. (2023) showing altitude (m.a.s.l.), mean annual precipitations (MAP), mean annual temperature (MAT), and distance between sampled sites (left). Principal component analysis (PCA) conducted upon the scaled environmental metadata across sites (right). OM: organic matter, EC: electric conductivity. **B.** Relative abundance of taxonomic orders (left). Taxa with abundance < 3% in all sites were merged into ‘other unabundant orders’. **C.** Relative abundance of most dissimilar functional categories between sites following the KEGG pathway annotations (left). Ranked functions contributing up to 10% of the cumulative overall dissimilarity according to a Similarity percentage analysis (SIMPER) are shown. Importance decreases from left to right. **B-C (right).** Principal coordinates analysis (PCoA) conducted upon the Hellinger-transformed abundances for taxonomic orders (n=72) and KEGG functional categories (n=491).

To understand how prokaryotic communities are taxonomically and functionally configured as physicochemical conditions change along the TLT, 17 nutrients were measured *in situ*. The geography and nutrient measurements of the six studied sites have been previously described Andreani-Gerard et al. (2024) (Fig. 1A). Metagenomic sequencing was performed at the sampled six sites, enabling us to generate taxonomic profiles and functional descriptions of the annotated genes for each metagenome, and to reconstruct collections of metagenome-assembled genomes (MAGs) at each site (see Methods). Physicochemical characterization evidenced the overall soil dissimilarity across the six sites (Fig. 1A and Table S1). The first discriminative principal component is driven by both macro (P, S, K, Mg, and Ca) and micro (Mn, Na, Fe, and B) nutrients, which separates S6 from the rest of sites. The second principal component orders the other five ecosystems according to environmental nitrogen (N), organic matter (OM), and two heavy metals (Cu and Zn). A clear separation of the sites was observed when omitting the sample from the Lejía lagoon; where the puna ecosystem (S2) exhibits a positive discrimination given by OM, Fe, Cu and Zn (Fig. S1 and Suppl. Text S1).

Alpha-diversity analyses included 146 million reads with assigned taxa (45% of the total), which were classified in 335 described prokaryotic OTUs (330 bacterial and 5 archaeal defined at 96.5% marker genes identity, Table S3). Increased richness was observed in sites S1 and S2, with 120 and 152 OTUs, Shannon’s indexes of 4.6 and 4.8, and Fisher’s log-series of 7.9 and 10.1, respectively, compared to the rest of sites (S3 to S6, avg: 40±10.6, 3.5±0.27, and 2.4±0.69). These results point to the prepuna and puna soil microbes as markedly more diverse than the sampled steppe communities, an observation that was previously reported for the prepuna (Mandakovic et al., 2023) and could relate to the semiarid vegetation belt that characterizes the puna (Frugone-Álvarez et al., 2023).

While the six sites were overall dominated by Actinobacteriota and Proteobacteria in line with previous surveys of desertic soil (Vásquez-Dean et al., 2020; Feng et al., 2020; Naidoo et al., 2022), taxonomic profiling revealed different microbial composition across sites even at the phylum, class and family ranks (Suppl. Text S2 and Fig. S2). Examination down to the rank of orders revealed little overlap of taxa between the surveyed ecosystems (Fig. 1B, Table S4), as confirmed by beta-diversity analyses separating the prepuna and puna from remaining sites along the PCoA’s second coordinate while the first coordinate strongly separates the Lejía lagoon’s community (S6) at all taxonomic ranks (Fig. 1B and Fig. S2).

To assess the impact of the observed taxonomic divergence on biological functions, we analyzed gene annotations from the PFAM, COG, KEGG pathways, and CAZyme databases (Fig. 1C and Fig. S3). A total of 8605, 4491, 429, and 125 entries were identified across the six samples, respectively. In average, 6876 (80%), 4070 (91%), 413 (96%), and 101 (81%) different functional categories from the respective nomenclatures were detected per sample (Table S5). Dissimilar functional categories were ranked using pairwise comparisons of Hellinger-transformed gene abundances through SIMPER analysis (see Methods). Functional profiles are detailed in Suppl. Text S3. Prepuna (S1) and puna (S2) were found to be enriched in genes associated to degradation of aromatic compounds like benzoate, whilst Lejía lagoon (S6) was characterized by motility associated functions. Sites S3, S4, and S5 showed no enrichment of the top KEGG functions contributing up to 10% of cumulative dissimilarity (Fig. 1C). PFAM analyses of the steppe microbiomes displayed an enrichment of several categories related to mobile genetic elements (e.g., transposases and phage integrases) and DNA repair (Fig. S3A). We observed that the most dissimilar COG annotation was an extracytoplasmatic receptor characterizing S2 and related to the uptake of tricarboxylates (TctC, Fig. S3B). Carbon metabolism was surveyed through the annotation of carbohydrate-active enzymes (CAZymes) which highlighted enrichment of glycosyl hydrolases (GH29, GH95, and GH3) in S6 and a broad glycosyl transferase (GT4) in S1 to S5 (Fig. S3C).

Overall, functional annotations, taxonomic profiles, and physicochemical characteristics, all confirm the heterogeneity of the TLT microbiomes, separating the puna and prepuna from the steppe samples and the lagoon (see Suppl. Text S4). These observations motivate a deeper exploration of the metabolism through dedicated models in order to suggest mechanistic hypotheses on the transect’s diversity.

### A systems biology strategy to simulate the metabolic potential according to nutrient environmental availability

Given the strong relationship revealed in the taxonomic and functional analyses of the metagenomic samples from the TLT, as well as the evidence of relationships with their physicochemical environment, we simulated the *metabolic potential* of the communities as a proxy of how they are prepared to respond to environmental nutritional shifts. We built a systems biology-based strategy that relies on metabolic modeling as the basis of a dynamical system for simulation. For this, a gene catalog was reconstructed for each metagenome while 120 high-quality MAGs, ranging from 5 to 44 per site and accounting in average for 15.1%±4.7% of relative abundance when mapped against metagenomic reads, were obtained across the six sites (Table S6). The originality of the approach lies in the construction of distinct and complementary models for each site, using their whole metagenome and their set of reconstructed MAGs. Metagenome-scale models (*MetaG-GEMs*) provide a global description of the metabolism of the entire community, independent of the taxa performing the functions. Building models based on MAGs (*MAG-GEM*) per site enables the study of putative populations that, although representing only a fraction of all prokaryotic species inhabiting the TLT, are abundant enough to be captured by genomic assemblies. Thus, the MAG-scale approach can be understood as a compromise that provides taxonomic identity to a subset of functions that are likely to be important in the corresponding communities, at the cost of possibly losing reactions from contigs that could not be binned. Overall, the objective of our strategy based on explainable models is to identify metabolic drivers associated with each site, and to determine taxa associated to their synthesis.

The first part of the method consisted in constructing two dynamical systems for each site using MAG-GEMs and MetaG-GEMs (Fig. 2A). The formalism used in the dynamical system is the one of metabolic modeling associated to a Boolean semantic. For the two systems associated to a site, we relied on a metabolic network reconstructed from its gene catalog (MetaG-GEM) and a collection of metabolic networks reconstructed for its MAGs (MAG-GEM). The choice of the Boolean semantics is motivated by our objective of providing a global description of the functions carried out by soil microorganisms. Metabolic networks yielded, in average, 5828±234.3 and 3774±583.5 unique reactions per sample from the assembled metagenomes and MAGs, respectively, of which more than 80% were gene-related (Table S7).

**Figure 2:**
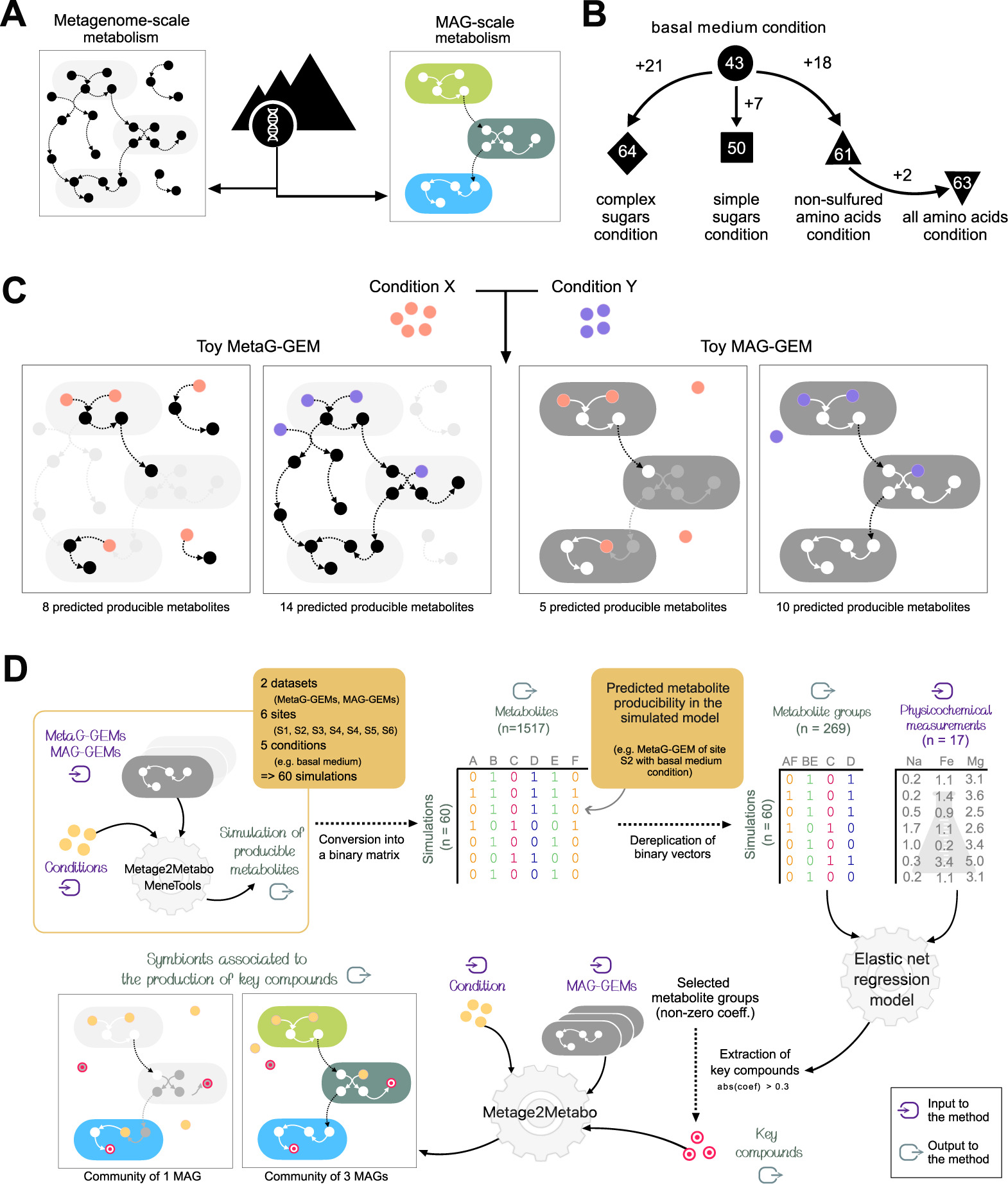
Overview of the metabolic modeling framework. **A.** Reconstruction of metabolic networks from sequence data for metagenomic (gene catalogs by site, left) and genomic (MAGs, right) datasets. Nodes are metabolites and arrows are reactions. Each sample’s metabolism can be abstracted as a MetaG-GEM built from the community-wide metabolic network, or a MAG-GEM obtained from the collection of metabolic networks resulting from each site’s MAGs. **B.** Definition of conditions to be simulated (user-provided seeds, Table 1 and Table S8). Numbers indicate the number of seed metabolites for each condition. **C.** Toy example of network expansion using a Meta-GEM (left) or a MAG-GEM (right). Pink and purple circles denotes two different conditions, represented as different available compounds. Black (resp. white) arrows and circles denote the reached metabolites by condition in the MetaG-GEM and MAG-GEM, respectively. **D.** Summary of the bioinformatics pipeline used in this work. Sixty predictions of reachable metabolites (scope) are performed with either MAGor MetaG-GEMs and five conditions for all six sites. Producible metabolites across conditions are summarized into a binary matrix. Matrix columns are dereplicated into groups of metabolites with identical producibility profiles across conditions. The latter are used as explanatory variables in the elastic net regression that aims at explaining the soil physicochemical metadata. Metabolite groups significantly associated (coefficient *>* 0.3) with environmental data are set as targets for the selection of MAG-GEMs minimal communities: key species (colored organisms) are involved in the biosynthesis throughout individual genomic capabilities (left, a single organism can produce the key compounds) and/or putative cross-feeding interactions (right, several interacting organisms are necessary).

The second step of the method consisted in defining simulation conditions for the dynamical systems. Considering that the Atacama Desert exhibits oligotrophic conditions and, thus, organic carbon and nitrogen supplies are expected to be scarce, we provided five scenarios simulating different nutritional sources to our concentration-independent Boolean approach (Fig. 2B). A *basal medium* is defined to contain inorganic compounds, a limited set of cofactors, coenzymes, and metal ions, with CO_2_ and HCO_3_ as inorganic carbon sources. The next two nutritional conditions were constructed to explore the effect of adding organic carbon, specifically, carbohydrates that enter glycolysis directly (*simple sugars*, Wang et al. (2019)), and compounds commonly found in soils with higher molecular weights, or that comprise alternative carbon sources (*complex sugars*, Swenson et al. (2015)). Finally, to assess the effect of supplying organic nitrogen, we provided a condition with all genomically-encoded amino acids (*all amino acids*) and another one excluding cysteine and methionine (*non-sulfured amino acids*) (see Methods, Table 1 and Table S8).

The third step consisted in running the simulations and computing the *metabolic potential*, the response of the dynamical system to the simulated conditions, described as sets of metabolites predicted to be producible. We illustrate in Figure 2C the impact of two initial conditions on the production of metabolites in a toy MetaG-GEM and its corresponding toy MAG-GEM. We observe that functions present in the metagenome but absent from MAGs may alter the set of producible metabolites, and that therefore the metabolic potential can vary with simulated conditions (Fig. 2C). In total, using the data from the TLT, 60 simulations were conducted, accounting for the five conditions and the two systems for each of the six sites (Fig. 2D).

The fourth step of the approach involved statistical analyses and additional simulations that aim at identifying relevant metabolites associated with each site, together with the taxa responsible for their production. The global framework of simulations and analyses is illustrated in Fig. 2D. Briefly, results of dynamical systems simulations were dereplicated in order to identify *metabolite groups* with similar producibility behaviors across simulations. Associations between these groups and physicochemical measurements of sites are identified using regression models. Metabolite groups with the strongest associations to measurements are referred to as key compounds and used in further simulations to predict MAGs associated to their production in each site. We detail in the following sections the outcomes of this systems biology framework.

### Metabolic modeling of soil prokaryotes outlines the potential of communities to adapt in demanding environments

Results of the simulations are presented in Fig. 3 and Fig. S4. Across sites, 1,517 unique metabolites were predicted to be producible in MetaG-GEMs, of which 1,166 (76.9%) were captured in MAG-GEMs. Producible metabolites ranged on average from 1,136.3±47.9 with the *basal medium* condition to 1,306.2±64.9 with *all amino acids* in MetaG-GEMs, and from 684.8±88.9 to 773.3±100.3 in MAG-GEMs, respectively (Fig. 3A). As expected, the MAGGEMs - that encompass a reduced portion of the microbiome - exhibit smaller scopes than those obtained through the MetaG-GEMs, regardless of the simulated conditions (Fig. 3B,C). The number of producible metabolites depends on the simulated condition. The highest number of producible metabolites in MAG-GEMs is exhibited by *complex sugars* and *all amino acids* conditions, and the latter presents the largest metabolic potential in MetaG-GEMs. This increased metabolic potential in *all amino acids* condition for both MAG- and MetaG-GEMs suggests that the addition of organic sulfur, i.e., cysteine and methionine, has a strong effect compared to solely organic nitrogen simulated in the *non-sulfured amino acids* condition. Additionally, puna (S2) reached the largest metabolic potential regardless of the conditions in the MetaG-GEMs, while the prepuna (S1) and one of the steppes (S4) consistently exhibited the lowest number of predicted metabolites in MAG- and MetaG-GEMs, respectively (Fig. 3B,C). Metabolites found to be producible in all 30 possible combinations of sites and conditions (core metabolites) prevail in both datasets, comprising 61.2% and 40.5% of all compounds in MetaG-GEMS (n=928) and MAG-GEMs (n=472), respectively (Table S9).

**Figure 3:**
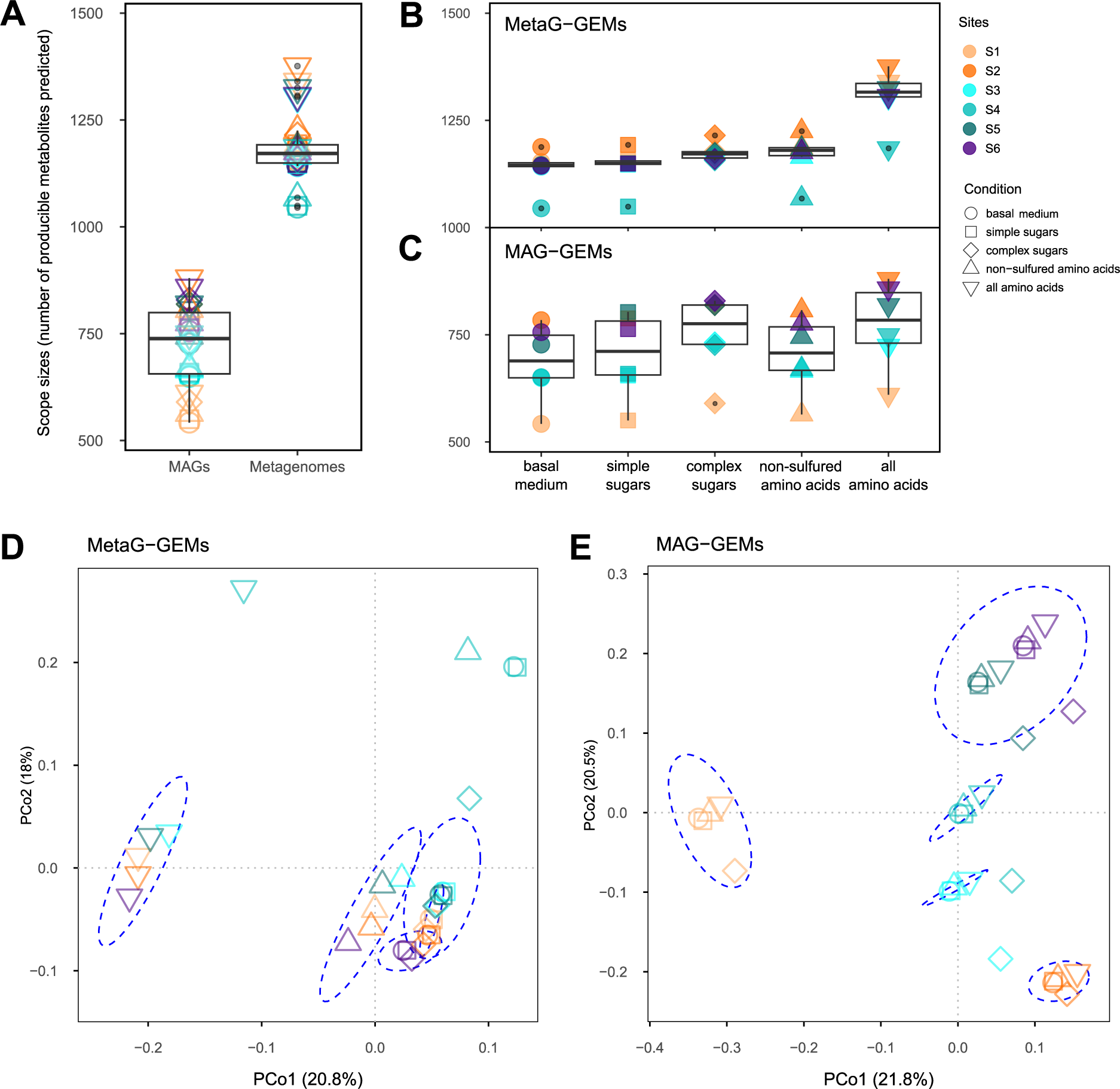
Quantitative description and ordination analysis of producible metabolites. **A.** Distribution of the number of producible metabolites across all 30 simulations (6 sites, 5 conditions) in MAG-GEMs and MetaG-GEMs. **B,C.** Number of producible metabolites according to the simulated conditions in MetaG-GEMs (B) and MAG-GEMs (C). **D,E.** PCoA obtained from the presence/absence matrix of producible compounds for MetaG-GEMs (D) and MAG-GEMs (E).

An ordination analysis was performed to compare producible metabolites across sites and conditions (Fig. 3D and 3E). Results show that MetaG- and MAG-GEMs exhibit different metabolic profiles. On one hand, the high proportion of core metabolites predicted from the MetaG-GEMs determines the overlap of all sites but S4 (see Suppl. Text S3) when carbon is provided with *basal medium*, *simple sugars*, and *complex sugars*. While profiles of reachable metabolites are more dissimilar in the *all* and *non-sulfured amino acids* conditions, the same five sites still cluster together. This underlines the strong impact of organic sulfur when compared to organic nitrogen and organic carbon, already implied by Fig. 3B. The effect of conditions on the simulated metabolic profiles, rather than the effect of sites themselves, suggests high functional redundancy across sites (Fig. 3D). On the other hand, most metabolic profiles of MAG-GEMs cluster by site (Fig. 3E) indicating that the subset of MAGs from each site harbor different metabolic pathways, specifically fitted to the physicochemical properties and nutrient contents along the TLT. These results suggest that microbial communities as a whole could act as a reservoir of functions with little variability across sites (Fig. 3D), whereas the most abundant players in each community, retrieved as MAGs, exhibit higher differences in their metabolic responses across sites (Fig. 3E).

### Selection of key metabolites driven by the environment

The binary matrix of metabolites (n = 1,517 unique metabolites) predicted to be produced across simulated conditions (Fig. 2D) was dereplicated by grouping metabolites with identical producibility patterns across all 60 simulations, raising 269 *metabolite groups*. Among those, 45% (n=121) harbored a unique metabolite, the median and average size of a metabolite group being 2 and 5.6, respectively; and the largest group, the one gathering core metabolites across datasets, contained 469 metabolites. Hierarchical clustering of metabolite group’s producibility vectors across sites and conditions (Fig. S5) highlights two main clusters: 211 metabolite groups that exhibited varying producibility status across sites but were rather insensitive to the simulated conditions (Fig. S5A), and 58 groups whose producibility was mostly driven by the amino acid-containing conditions while being overall insensitive to sites (Fig. S5B). In the latter cluster 16 metabolite groups associated with *non-sulfured amino acids* and 42 to the inclusion of cysteine and methionine. We refer to interactive files in Supplementary material for a detailed description of biochemical families that distinguish sites across simulations in MAG-GEMs and MetaG-GEMs. Results highlight missing functions related to biosynthesis of polysaccharides, glycoconjugates, terpenoids, and of various lipids in the MetaG-GEM from S4 with respect to remaining sites. In MAG-GEMs, those functions were only present for the puna (S2).

We used a regression model to associate the *in situ* physicochemical measurements (n = 17, Fig. 1A) with metabolite groups extracted from simulations (Fig. 2D). Our results show that 82 out of the 269 unique metabolite groups (30.1%) had nonzero coefficients in the model; meaning that 292 out of the 1,517 unique metabolites (19.3%) were fitted as relevant for at least one environmental variable (Table S10). Since coefficient values accounts for the strength and direction of the corresponding relationship, we focused on metabolite groups with absolute values greater than 0.3 (n=39), obtaining 171 metabolites which we refer to as *key metabolites*. A hierarchical clustering of the selected metabolite groups revealed three main clusters of environmental variables (EV1, EV2, and EV3, Fig. 4A). Namely, EV1 is defined by K, Na, Fe, P, Mn, Mg, S, B, Ca, and electric conductivity; EV2 by organic matter (OM), Zn, and N; and EV3 by pH, Cu, NO_3_ and NH_4_. Then, looking at the distribution of the in situ measurements across sites (Fig. 4B), we observed that EV1 and EV2 mainly cluster variables for which the Lejía lagoon (S6) and the puna (S2) microbiomes are outliers respectively, implying that they likely constitute major abiotic stressors for microorganisms inhabiting these rare ecosystems, whereas EV3 has no evident relation to geography.

**Figure 4:**
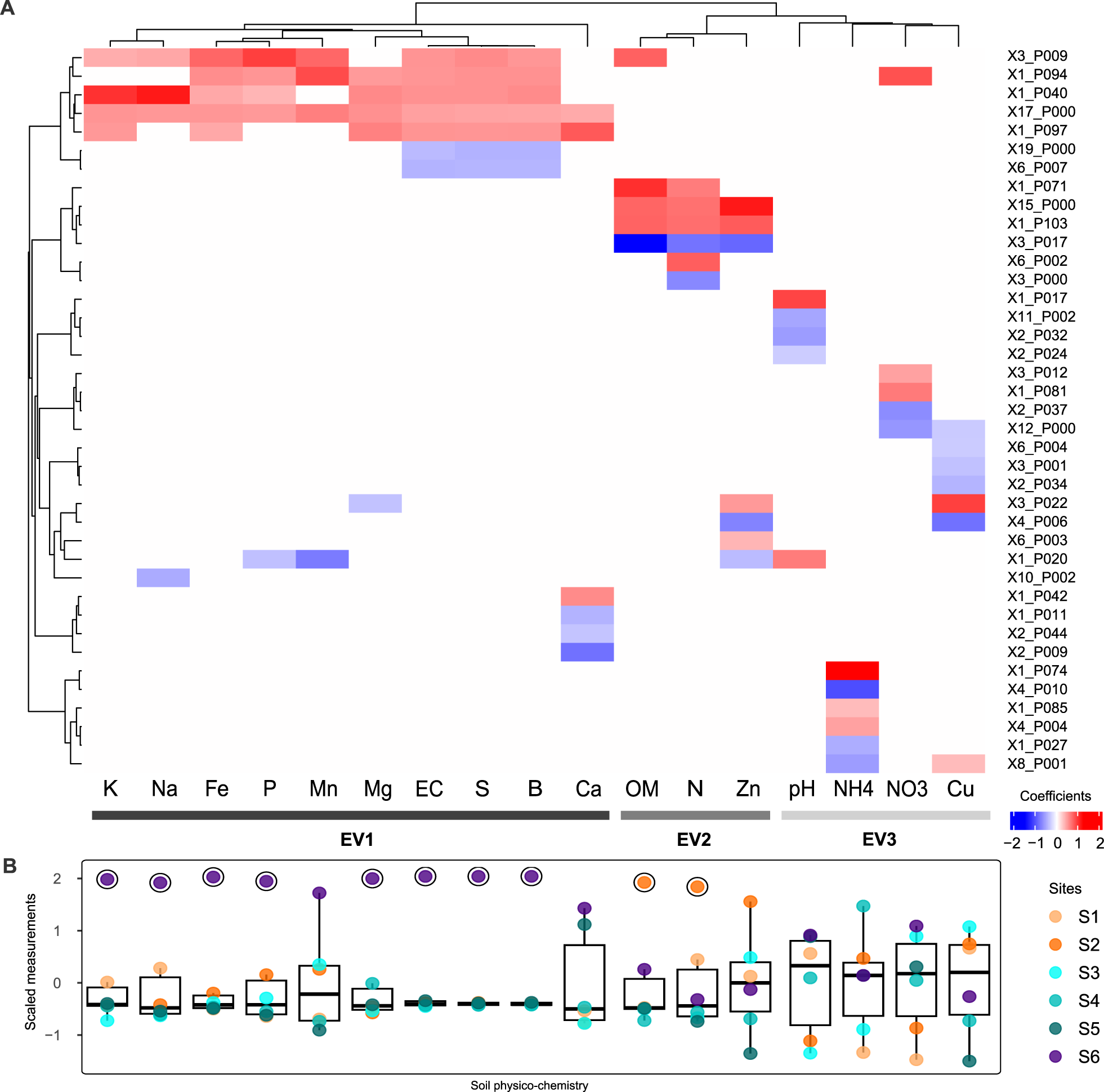
Key metabolite groups proposed to be driven by the environmental pressures of the TLT. **A.** Heatmap of metabolite groups selected with an elastic net regression model as best predictors for explaining environmental metadata. The first term of IDs indicates the number of metabolites in each group. Data is shown if the absolute value of the fitted coefficient was *≥*0.3 (see Table S10). A total of 171 key metabolites emerged from these 39 metabolite groups (rows). Hierarchical clustering of columns enabled three groups of environmental variables (EVs) to arise. **B.** Boxplot show the scaled values of *in situ* soil physicochemical measurements comprising the environmental metadata. Outliers are displayed inside black circles. OM: organic matter, EC: electric conductivity.

We further surveyed the link between sites and obtained metabolite groups associated with the first two clusters of environmental variables. Firstly, EV1 was positively associated to five metabolite groups representing 23 metabolites linked to nitrogenated pathways (e.g., Lleucine, L-valine, L-arginine, N-acetylneuraminate, and N-acetylmannosamine degradation, Lisoleucine biosynthesis) and osmotic stress (see Suppl. Text S5). Secondly, EV2 was positively associated to 2 metabolite groups representing 23 metabolites mostly linked to carbon cycling (e.g., staurosporine, violacein, and flavonoid biosynthesis, and aromatic degradation). This observation is consistent with, and further expands results of the functional profiling above, showing enrichment in some functions related to carbon catabolism and transport in the puna (Fig. 1C and Suppl. Text S6). For a description of structural ontology of key compounds from the puna (EV2) and the lagoon (EV1), we encourage the interested reader to navigate the interactive html files provided in the Supplementary material.

### Identification of MAGs involved in the production of key metabolites

To identify the genomes having the potential to produce the 171 key metabolites associated with the sites’ physicochemical measurements, we set up a new simulation using the MAG-GEM of each site with the *basal medium* condition as inputs to Metage2Metabo, which predicts minimal-size communities (Figs. 2D and 5A, see Methods). This reverse engineering approach aims at capturing a refined signal associating key metabolites to key species able to produce them. A first observation was that 90 key compounds were produced only in MetaG-GEMs simulations (Fig. 5B). Although precise information on the taxa responsible for their production cannot be inferred, these 90 metabolites constitute the environmentally-driven signal for whole communities. Hence, despite the MetaG-GEMs demonstrate a high functional overlap, as pointed out in Section *Metabolic network reconstruction and modeling*, this method can pinpoint metabolic functions that are specifically related to geography and could not be captured from the first set of simulations (Fig. 3D). We therefore assessed the producibility of the remaining 81 key metabolites (27 metabolite groups) in the six sites (Fig. 5C).

**Figure 5:**
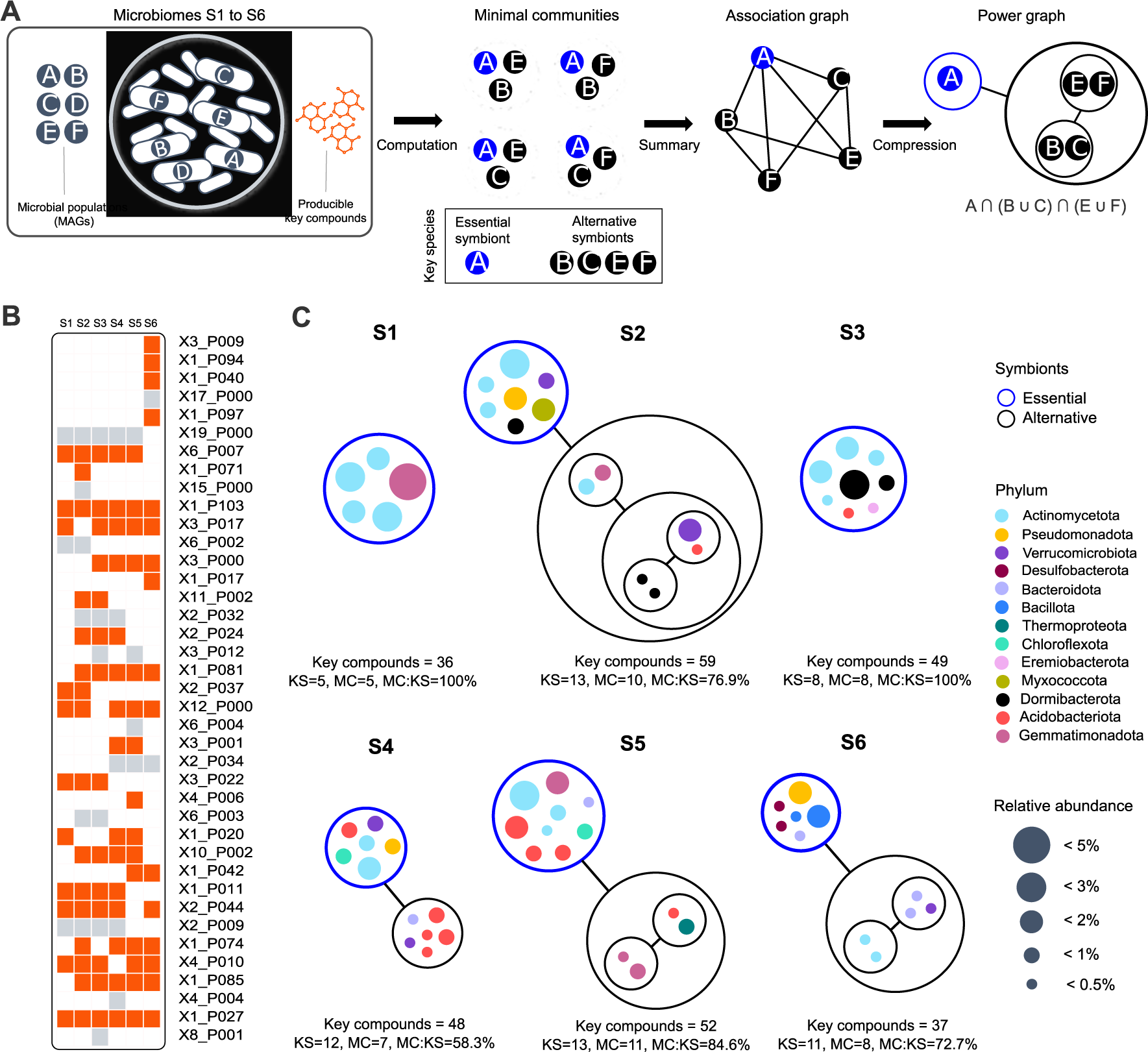
Minimal communities potentially able to produce key metabolites. **A.** Illustration of the reverse engineering simulation protocol for identification of MAGs involved in the biosynthesis of key compounds with Metage2Metabo (Belcour et al., 2020), see also Fig. 2D. **B.** Panel showing the producibility of metabolite groups (n=39) containing key compounds (n=171) by site (see Table S11). Orange squares depict metabolite groups (n=27) containing key compounds (n=81) that are producible in MAG-GEMs. Gray squares depict metabolite groups (n=12) containing key compounds (n=90) that were only producible in MetaG-GEMs and, thus, had no effect in the computation of minimal communities, which is only possible for the genome-resolved approach. The first term of IDs indicate the number of key compounds in each metabolite group. **C.** Power graphs summarizing the structure of all predicted minimal communities per site. Blue and black circles should be interpreted as “AND” and “OR”, respectively, meaning that all MAGs inside blue circles are required for the production of the key compounds provided as targets (essential symbionts) whereas only one MAG is needed inside each black circle (alternative symbionts). Lines represent the sequence of decision-making for the combinatorics of possible solutions and should be interpreted as “AND”. Relative abundance represent the percentage of metagenomic reads mapping to each MAG, i.e., populations in panel (A). KS: key species, MC: minimal community.

Our findings show that, out of the 120 MAGs, 62 were involved in the biosynthesis of at least one key metabolite. From these, 41 MAGs were essential symbionts (Fig. 5C), meaning that they occurred in all predicted minimal communities and, thus, may constitute load points for the producibility of the key metabolites (Tables S6 and S11). This suggests that one third of the reconstructed MAGs carry unique metabolic machinery related to at least one key metabolite. In average, MAGs were predicted to produce 46.8±8.9 key metabolites by site, of which 26.3±6.1 were predicted to be producible only through cooperative mechanisms with other MAGs from the site (Table S11).

We dived further into the structure of communities per site by enumerating all possible minimal communities associated with key metabolite biosynthesis and analyzing the co-occurrence patterns regarding included MAGs and their combinations (see Methods). MAGs occurring in at least one minimal community at a specific site are referred to as *key species* (Belcour et al., 2020), denoting the functional redundancy of the microbiome with respect to the production of targeted metabolites when the number of key species is larger than the size of the minimal community. In other words, some solutions may predict one key species over another without altering the ability for the minimal community to produce the targets. In our case, the predicted minimal communities exhibited different microbial structures regarding the number, abundance, and taxonomy of MAGs involved in the production of key metabolites (Fig. 5D). Some sites (S1, S3) led to rigid communities: an unique community was predicted as able to sustain the producibility of key metabolites, meaning that no MAG could be removed or replaced while preserving the community’s biosynthesis potential (MC:KS=100%, MC: minimal community, KS: key species). These theoretical abstractions indicate more fragile communities, where any environmental perturbation could potentially compromise the ability of the community to sustain the biosynthesis of such metabolites (Xun et al., 2021). On the opposite side, we found a range of more complex structures providing many alternative community compositions for a single function, such as for S4 (MC:KS=58.3%), to a couple of replacements for a few functions like in S2, S5, and S6 (MC:KS>70%). These alternative symbionts increase the combinatorics of possible community assemblies a soil microbiome can resort to in order to reach the targeted metabolic products. In these cases, metabolic redundancy suggests robustness to changes in community composition upon environmental circumstances (Shade et al., 2012). Such reserve of functions offers plasticity and, ultimately, resilience to the whole communities where these MAGs were recovered from (Allison and Martiny, 2008).

We finally surveyed the abundance of the MAGs proposed as key species in the metagenomic datasets. On average, key species accounted for 8.8% of total relative abundance per site. S6 behaved as an outlier since selected MAGs accounted for 3.5% only of the total metagenomic abundance, which suggests that microbes sampled from the Lejía lagoon probably have a complex underlying structure that our approach overlooked because of assembly limitations. Nevertheless, this result also implies that unabundant prokaryotes are likely to catalyze critical metabolic steps in ecosystem functioning, as argued by Wang et al. (2021). On the other hand, we observed that MAGs from S1, all defined as essential symbionts, encompassed the highest cumulative abundance of key species (13.1%) across the TLT ranging from 1.6% to 4.7% per MAG, compared to an average of 0.8% for essential symbionts in the rest of sites. This observation suggests that the metabolic response of microbial communities may fall on a few abundant members, likely due to the successful strategies they employ in the demanding environmental conditions of the TLT that severely limit other microbes to thrive (Stone et al., 2021). Regarding taxonomy, we observed that symbionts of Actinomycetota and Acidobacterota constituted key species in every site, suggesting that these phyla harbour important functions in the microbiomes, and especially the former that is predicted in every enumerated community (Fig. 5D). For a complete taxonomic description of minimal communities, we refer the interested reader to Suppl. Text S7. Overall, this approach of community selection can pinpoint MAGs and taxa exhibiting important roles in the community regarding functions of interest. Combining this information to taxonomy and abundance data can raise hypotheses on the global organization of the microbiome.

## Discussion

### A functional description of the TLT using metabolic modeling

In this work, we characterize the metabolic potential of six contrasting microbiomes from an altitudinal gradient in the Atacama Desert, and explore their relationship with soil physicochemical metadata by using a systems biology strategy. In general, our results show that the addition of organic carbon in the forms of simple and complex sugars do not signify a major change in the predicted scopes of the TLT microbiome when compared to simulations performed only with inorganic carbon sources. Although carbon fixing members are known to be scarce in soils (Garritano et al., 2022), our abundance-independent approach underscores the ability of those few members to contribute to carbon uptake for the whole community to which they belong and the underlying interactions of cooperation. We also observed an increased sensitivity towards the input of organic sulfur, in contrast to organic nitrogen, that highlights the relevance of methionine and cysteine as precursors in metabolite biosynthesis. Methionine and cysteine are the only sulfur-containing amino acids incorporated into proteins. The former, in the form of N-formylmethionine, initiates the synthesis of nearly all prokaryotic proteins, while S-adenosylmethionine, a highly versatile cofactor, is involved in methyl groups and 5’-deoxyadenosyl group transfers, polyamine synthesis, and numerous other processes. In contrast, cysteine forms disulfide bonds that determine protein structure and participate in protein-folding pathways (Brosnan and Brosnan, 2006).

Given that key metabolites comprised a small fraction of the simulated scopes, we were able to sharpen the differences between sites retrieved via metabolic modeling and further link them with seventeen environmental drivers. On one side, the capabilities for pigment and antioxidant biosynthesis and for aromatic degradation of the puna microbiome were related to the high contents of organic matter, nitrogen, and zinc in that area. A study assessing the functional potential of soil microbes associated to transitional hygrophilous plants from the Atacama Salar showed an enrichment of certain metabolic pathways for the degradation of organic matter and aromatic compounds and the biosynthesis of amino groups (Ramos-Tapia et al., 2022). We propose that the distinctive vegetation belt that characterizes the puna ecosystem (Dussarrat and et al, 2025) is a critical source of nitrogen throughout the exudation of organic compounds into the soil that also chelate zinc, immobilizing it for slow release. This would significantly improve carbon and nitrogen availability in an ecosystem adapted to extreme nutrient limitations (Vikram et al., 2016).

On the other side, several alternative pathways for amino acid biosynthesis and degradation detected at the Lejía lagoon’s shore were driven by abiotic factors related to salinity, electric conductivity, and sulfur, among others. The chemical structures of some of these nitrogenated compounds classified them as short chain fatty acids and glycans, suggesting that they could contribute to the metabolic response to temporal shifts in water availability by improving water retention and serving as energy reservoir (Lennon and Jones, 2011; Dinnbier et al., 1988). These results complement our recent work assessing specialized metabolism on the same soil samples that highlighted some bacterial members of the lagoon microbiome with niche-adaptations for the acquisition of organic nitrogen throughout heterocyst glycolipid-like mechanisms and for osmotic stress resistance throughout ectoines (Andreani-Gerard et al., 2024).

### MAGs highlight emergent properties from functionally redundant gene reservoirs

The metabolic potential recovered from the TLT metagenomes evidenced a high functional redundancy across the sampled ecosystems, where more than 60% of producible metabolites predicted with the community-wide approach were found in every site regardless of the simulation conditions. This is consistent with previous observations made on 845 soil communities across 17 climate zones around the globe where, due to functional redundancy, microbial functions based on gene abundances were more stable regarding geography than taxonomy and soil properties (Chen et al., 2022). Another effort surveying 962 metagenomic studies from nine different ecosystems showed that, while metabolic overlap in soils is overall lower compared to other environments (e.g., marine and freshwater), extreme environments exhibited the highest functional redundancy (Hester et al., 2019). Given that functional redundancy is a necessary condition for taxonomic turnover within functional groups (organisms capable of performing a specific metabolic function) (Louca et al., 2018), we hypothesize that the microbial communities inhabiting the TLT depend on a fairly exhaustive *gene reservoir* to withstand the environmental perturbations they may sporadically encounter in such extreme conditions. Nevertheless, our regression-based strategy distinguished 90 compounds that accounted for the metabolic differences – at first sight hidden behind such redundancy – between sites, when whole communities were assessed.

In this sense, it has been speculated that populations with similar metabolic repertoires may specialize on distinct nutrients and, thus, express separate ‘realized’ niches rather than ‘fundamental’ niches at the transcriptional level (Hutchinson, 1957; Louca et al., 2018). Considering that the metabolic functions performed by a given population are finely tuned by the environment, including the presence and activity of other community members, only a few members of a same functional group may emerge to actively perform a given function. Thus, while some members can exhibit alternative modes to gain energy, others may simply be inactive due to differing enzyme efficiencies, growth kinetics, and other traits influencing their growth rates under specific conditions (Louca et al., 2018). The latter would explain why MAGs in our dataset display a site-driven metabolic behavior fitted to their corresponding geographies, with little functional overlaps. Therefore, we propose that the genome-resolved approach offers valuable insights into the emergent properties of metabolisms thriving in specific contexts and that reconstructing MAG-scale metabolic networks is particularly useful for identifying adaptations that enable organisms to dominate a community under defined environmental conditions.

Assessment of MAGs enabled us to identify key species involved in biosynthesis of key metabolites, this is, showing an association with the environment. These key species are classified as essential or alternative symbionts depending on whether they were strictly required for the production of targeted metabolites or whether they could be functionally replaced by another member of the community (Belcour et al., 2020). Although it is uncertain whether MAGs herein defined as essential symbionts exert a critical role in organizing the structure of their respective soil microbiomes, they exhibit unique metabolic functions that connect tightly with biosynthetic steps found in the genomes of other community members, *i.e.*, metabolic handoffs (Anantharaman et al., 2016; Hug and Co, 2018), and, hence, could constitute keystone taxa (Banerjee et al., 2018). We hypothesize that some of these metabolic dependencies may constitute cross-feeding through putative cooperation events and that the removal of the proposed keystone taxa could alter microbiome stability, potentially causing downstream impacts on ecosystem processes (Mataigne et al., 2021). Finally, we acknowledge that our approach may have overlooked other organisms displaying keystone roles due to limitations in the genomic assemblies related to low abundance (Ejaz et al., 2024) and that, even though minimal communities is a reasonable mathematical solution for delving into ecological functioning of microbiomes, it may not comprehensively reflect the (non-minimal) mechanisms employed in nature.

### The potential of metabolic modeling to decipher community-wide and genome-resolved functions

A methodological contribution of this work is the systems biology framework that relies on dynamical systems for modeling the metabolic potential of metagenomes and collections of MAGs in the Atacama soils. The use of numerical models such as flux balance analysis (reviewed in Cerk et al. (2024)) would be hardly applicable in a context where metabolic networks were reconstructed automatically and curation is limited for non-model organisms of such extreme environment. Dynamical system simulations with ordinary differential equations on the other hand would require setting up many parameters. Here, the discrete model of metabolic producibility is an approximation of such numerical models that offers flexibility and predicts fixed points of the dynamical system (Frioux et al., 2020). As a model, it enables nonetheless to go beyond pathway description because it predicts the effect of environmental nutrients on the metabolism. The approach used here relies on the network expansion algorithm (Ebenhöh et al., 2004) which was implemented in a logic paradigm and extended to the ecosystem level (Frioux et al., 2018; Belcour et al., 2020).

Our model uses both community-wide metabolic networks, constructed from gene catalogs of metagenome assemblies by site, and genome-scale metabolic networks. The former (MetaG-GEMs) captures the most functions of the ecosystem as it takes into account any gene that is assembled, but it does not provide information in regards to which organism carries each function. The latter (MAG-GEMs) on the other hand enables predicting the relationship between taxonomy and function, although it underestimates the complexity of the community due to the difficulty of MAG reconstruction in complex microbiomes such as those from soils (Ejaz et al., 2024). Improvements in genome-resolved metagenomics, for instance with long-read sequencing will increase the number of genomes that can be obtained, and thereby enhance the quality of associated metabolic models (Cerk et al., 2024). Each of the six sites of the transect in this study is analyzed both at the level of the whole community’s metabolic network or with its associated collection of MAG-derived networks. In both cases, our approach highlights the sensitivity of the metabolism to varying nutrients sources and a core set of functions that could be reachable regardless of nutrients available in the environment.

Among the simplifications performed by the model, any metabolite producible by a taxon will be considered available to the community, hence the cost of transporting metabolites is ignored. While this characteristic could raise false positive cross-feeding predictions, it arises from both a modeling limitation related to transporter annotation (Casey et al., 2024) and ecological considerations that stress the importance of metabolic exchanges in microbiomes, and their expected limited cost for microbial fitness Pacheco et al. (2019). In addition to the above caveat, the cost of enzyme biosynthesis is not modeled, whereas it can have a strong impact on microbial adaptation to an environment with scarce nutrient availability (Noor et al., 2016; Goelzer et al., 2015; Domenzain et al., 2022). We further acknowledge that metabolic reactions databases are incomplete, and that many proteins remain of unknown function, suggesting that an important part of the transect metabolism still has to be elucidated.

## Conclusions

We conceived a generic modeling framework, suitable for non-model microorganisms and scalable for large datasets, that facilitates progress toward disentangling the complex metabolic interactions that shape microbiome functioning. With few inputs, i.e., sequence data, custom nutritional scenarios, and physicochemical metadata, we link metabolic insights with taxonomy and, more importantly, with the environment. Prediction of the metabolic potential under varying conditions, inference of key metabolites associated with environmental drivers, and identification of key species involved in the biosynthesis of those metabolites, allowed us to uncover niche adaptations evolved along the Talabre-Lejía transect, such as the influence of soil organic matter on aromatic degradation and of salinity and other stressful abiotic factors on nitrogen cycling. We also captured the crucial role of organic forms of sulfur in this oligotrophic environment that stood over the impact of carbon and nitrogen on microbial metabolism. Finally, our choice of modeling entire communities and individual organisms in parallel allowed to fathom the functional overlap often found in metagenomes as a gene reservoir that provides whole microbial communities means to adapt to future environmental shifts, MAGs accounting for single populations as divergent, emergent properties of microbiome functioning adapted to current environmental conditions, and the added value of cooperation for enduring them.

## Acknowledgements

CF and CM are supported by the French National Research Agency (ANR) France 2030 PEPR Agroécologie et Numérique MISTIC ANR-22-PEAE-0011. AM, AS, CA, CF, YL are supported by the Inria associated team “Symbiodiversity”. CM is supported by French Région Nouvelle Aquitaine.

AM, CA, NJ, RP were supported by the Center for Mathematical Modeling (CMM) BASAL fund FB210005 for center of excellence from ANID-Chile, Millennium Institute Center for Genome Regulation (Project ANID–MILENIO-ICN2021_044) and grant Exploración number 13220002.

Experiments presented in this paper were carried out using the supercomputing infrastructure of NLHPC at CMM, Universidad de Chile (see https://www.nlhpc.cl/infraestructura/), and the PlaFRIM experimental testbed, supported by Inria, CNRS (LABRI and IMB), Université de Bordeaux, Bordeaux INP and Conseil Régional d’Aquitaine (see https://www.plafrim.fr).

## Authors contributions

Conceptualization: AM, AS, CA, CF; Data curation: CA, CM, RP; Formal analysis: CA; Visualization: CA, CF; Funding acquisition: AM, AS, CF; Investigation: AM, AS, CA, CF, MG, NJ; Methodology: CA, CF, CM, PHG, RP, YLC; Writing - original draft: AM, AS, CA, CF; Writing - review & editing: AM, AS, CA, CF, MG, NJ, VC, YLC.

## Supplementary information

The manuscript includes three Supplementary Files: *supplementary_file1.xlsx* with Tables S1 to S11, *supplementary_file2.pdf* with Table S12, Figs. S1 to S5, and Supplementary Texts S1 to S7, and *supplementary_file3.zip* with the interactive htmls for visualizing structural ontology of key compounds.

## Data availability

Nucleotide sequences of metagenomes and MAGs are deposited at the NCBI database under the BioProject accession PRJNA1104199. Generation of genbank files from sequence data can be reproduced with the script deposited at https://github.com/rpalmavejares/meta_ gbk_generator. Metabolic networks are deposited at Zenodo 10.5281/zenodo.14537000. Statistical analyses and plots can be reproduced with the script deposited at https://github.com/cmandreani/models_Atacama.

## Copyright

A CC-BY public copyright license has been applied by the authors to the present document and will be applied to all subsequent versions up to the Author Accepted Manuscript arising from this submission, in accordance with the grant’s open access conditions.

## References

Aite, M., Chevallier, M., Frioux, C., Trottier, C., Got, J., Cortés, M. P., Mendoza, S. N., Carrier, G., Dameron, O., Guillaudeux, N., Latorre, M., Loira, N., Markov, G. V., Maass, A., and Siegel, A. (2018). Traceability, reproducibility and wiki-exploration for “à-la-carte” reconstructions of genome-scale metabolic models. PLOS Computational Biology, 14(5):1– 25.

Allison, S. D. and Martiny, J. B. H. (2008). Colloquium paper: resistance, resilience, and redundancy in microbial communities. Proc. Natl. Acad. Sci. U. S. A., 105 Suppl 1(supplement_1):11512–11519.

Alneberg, J., Bjarnason, B. S., de Bruijn, I., Schirmer, M., Quick, J., Ijaz, U. Z., Lahti, L., Loman, N. J., Andersson, A. F., and Quince, C. (2014). Binning metagenomic contigs by coverage and composition. Nat. Methods, 11(11):1144–1146.

Anantharaman, K., Brown, C. T., Hug, L. A., Sharon, I., Castelle, C. J., Probst1, A. J., Thomas1, B. C., Singh, A., Wilkins, M. J., Karaoz, U., Brodie, E. L., Williams, K. H., Hubbard, S. S., and Banfield, J. F. (2016). Thousands of microbial genomes shed light on interconnected biogeochemical processes in an aquifer system. Nat. Commun., 7(13219).

Andreani-Gerard, C. M., Cambiazo, V., and González, M. (2024). Biosynthetic gene clusters from the atacama desert. mSphere.

Arroyo, M. T. K., Squeo, F. A., Armesto, J. J., and Villagran, C. (1988). Effects of aridity on plant diversity in the northern chilean andes: Results of a natural experiment. Ann. Mo. Bot. Gard., 75(1):55.

Banerjee, S., Schlaeppi, K., and van der Heijden, M. G. (2018). Keystone taxa as drivers of microbiome structure and functioning.

Belcour, A., Frioux, C., Aite, M., Bretaudeau, A., Hildebrand, F., and Siegel, A. (2020). Metage2metabo, microbiota-scale metabolic complementarity for the identi1cation of key species. eLife.

Boon, E., Meehan, C. J., Whidden, C., Wong, D. H.-J., Langille, M. G. I., and Beiko, R. G. (2014). Interactions in the microbiome: communities of organisms and communities of genes. FEMS Microbiol. Rev., 38(1):90–118.

Brosnan, J. and Brosnan, M. (2006). The sulfur-containing amino acids: an overview. J Nutr, 136((6 Suppl)):1636S–1640S.

Buchfink, B., Xie, C., and Huson, D. H. (2014). Fast and sensitive protein alignment using DIAMOND. Nature Methods, 12(1):59—60.

Budinich, M., Bourdon, J., Larhlimi, A., and Eveillard, D. (2017). A multi-objective constraint-based approach for modeling genome-scale microbial ecosystems. PLoS ONE, 12.

Bushnell, B., Rood, J., and Singer, E. (2017). Bbmerge – accurate paired shotgun read merging via overlap. PLOS ONE, 12(10):1–15.

Casey, J., Bennion, B., D’haeseleer, P., Kimbrel, J., Marschmann, G., and Navid, A. (2024). Transporter annotations are holding up progress in metabolic modeling. Frontiers in Systems Biology, 4:1394084.

Cerk, K., Ugalde-Salas, P., Nedjad, C. G., Lecomte, M., Muller, C., Sherman, D. J., Hildebrand, F., Labarthe, S., and Frioux, C. (2024). Community-scale models of microbiomes: Articulating metabolic modelling and metagenome sequencing. Microbial Biotechnology, 17(1):e14396.

Chaumeil, P.-A., Mussig, A. J., Hugenholtz, P., and Parks, D. H. (2019). GTDB-Tk: a toolkit to classify genomes with the Genome Taxonomy Database. Bioinformatics, 36(6):1925–1927.

Chen, H., Ma, K., Lu, C., Fu, Q., Qiu, Y., Zhao, J., Huang, Y., Yang, Y., Schadt, C. W., and Chen, H. (2022). Functional redundancy in soil microbial community based on metagenomics across the globe. Frontiers in Microbiology, 13.

Cock, P. J. A., Antao, T., Chang, J. T., Chapman, B. A., Cox, C. J., Dalke, A., Friedberg, I., Hamelryck, T., Kauff, F., Wilczynski, B., and Hoon, M. J. L. d. (2009). Biopython: freely available Python tools for computational molecular biology and bioinformatics. Bioinformatics, 25(11):1422–1423.

Dinnbier, U., Limpinsel, E., Schmid, R., and Bakker, E. P. (1988). Transient accumulation of potassium glutamate and its replacement by trehalose during adaptation of growing cells of escherichia coli k-12 to elevated sodium chloride concentrations. Archives of Microbiology, 150:348–357.

Domenzain, I., Sánchez, B., Anton, M., Kerkhoven, E. J., Millán-Oropeza, A., Henry, C., Siewers, V., Morrissey, J. P., Sonnenschein, N., and Nielsen, J. (2022). Reconstruction of a catalogue of genome-scale metabolic models with enzymatic constraints using gecko 2.0. Nature communications, 13(1):3766.

Drula, E., Garron, M.-L., Dogan, S., Lombard, V., Henrissat, B., and Terrapon, N. (2022). The carbohydrate-active enzyme database: functions and literature. Nucleic Acids Res., 50(D1):D571–D577.

Dussarrat, T. and, et al (2025). Rhizochemistry and soil bacterial community are tailored to natural stress gradients. Soil Biology and Biochemistry.

Díaz, F. P., Frugone, M., Gutiérrez, R. A., and Latorre, C. (2016). Nitrogen cycling in an extreme hyperarid environment inferred from *δ*^15^N analyses of plants, soils and herbivore diet. Scientific Reports, 6(22226).

Ebenhöh, O., Handorf, T., and Heinrich, R. (2004). Structural analysis of expanding metabolic networks. Genome informatics. International Conference on Genome Informatics, 15(1):35– 45.

Ejaz, M. R., Badr, K., Hassan, Z. U., Al-Thani, R., and Jaoua, S. (2024). Metagenomic approaches and opportunities in arid soil research. Science of The Total Environment, 953.

Eshel, G., Araus, V., Undurraga, S., Soto, D., Moraga, C., Montecinos, A., Moyano, T., Maldonado, J., Díaz, F., Varala, K., Nelson, C., Contreras-Lóez, O., Pal-Gabor, H., Kraiser, T., Carrasco-Puga, G., Nilo-Polanco, R., Zegar, C., Orellana, A., Montecino, M., Maass, A., Allende, M., DeSalle, R., Stevenson, D., González, M., Latorre, C., Coruzzi, G., and Gutiérrez, R. (2021). Plant ecological genomics at the limits of life in the atacama desert. 118.

Feng, W., Zhang, Y., Yan, R., Lai, Z., Qin, S., Sun, Y., She, W., and Liu, Z. (2020). Dominant soil bacteria and their ecological attributes across the deserts in northern china. European Journal of Soil Science, 71:524–535.

Fierer, N. (2017). Embracing the unknown: disentangling the complexities of the soil microbiome. Nature Reviews Microbiology, 15(10):579–590.

Finn, R. D., Mistry, J., Schuster-Böckler, B., Griffiths-Jones, S., Hollich, V., Lassmann, T., Moxon, S., Marshall, M., Khanna, A., Durbin, R., Eddy, S. R., Sonnhammer, E. L. L., and Bateman, A. (2006). Pfam: clans, web tools and services. Nucleic Acids Res., 34(Database issue):D247–51.

Frioux, C., Dittami, S. M., and Siegel, A. (2020). Using automated reasoning to explore the metabolism of unconventional organisms: a first step to explore host–microbial interactions. Biochemical Society Transactions, 48(3):901–913.

Frioux, C., Fremy, E., Trottier, C., and Siegel, A. (2018). Scalable and exhaustive screening of metabolic functions carried out by microbial consortia. Bioinformatics, 34(17):i934–i943.

Frugone-Álvarez, M., Contreras, S., Meseguer-Ruiz, O., Tejos, E., Delgado-Huertas, A., ValeroGarcés, B., Díaz, F. P., Briceño, M., Bustos-Morales, M., and Latorre, C. (2023). Hydroclimate variations over the last 17,000 years as estimated by leaf waxes in rodent middens from the south-central atacama desert, chile. Quaternary Science Reviews, 311:108084.

Galperin, M. Y., Wolf, Y. I., Makarova, K. S., Vera Alvarez, R., Landsman, D., and Koonin, E. V. (2021). COG database update: focus on microbial diversity, model organisms, and widespread pathogens. Nucleic Acids Res., 49(D1):D274–D281.

Garritano, A. N., Song, W., and Thomas, T. (2022). Carbon fixation pathways across the bacterial and archaeal tree of life. PNAS Nexus, 1(5).

Goelzer, A., Muntel, J., Chubukov, V., Jules, M., Prestel, E., Nölker, R., Mariadassou, M., Aymerich, S., Hecker, M., Noirot, P., et al. (2015). Quantitative prediction of genome-wide resource allocation in bacteria. Metabolic engineering, 32:232–243.

Gu, Z. (2022). Complex heatmap visualization. iMeta.

Hester, E. R., Jetten, M. S., Welte, C. U., and Lücker, S. (2019). Metabolic overlap in environmentally diverse microbial communities. Frontiers in Genetics, 10.

Huerta-Cepas, J., Szklarczyk, D., Heller, D., Hernández-Plaza, A., Forslund, S. K., Cook, H., Mende, D. R., Letunic, I., Rattei, T., Jensen, L. J., von Mering, C., and Bork, P. (2019). eggNOG 5.0: a hierarchical, functionally and phylogenetically annotated orthology resource based on 5090 organisms and 2502 viruses. Nucleic Acids Res., 47(D1):D309–D314.

Huerta-Cepas, J., Szklarczyk, D., Heller, D., Hernández-Plaza, A., Forslund, S. K., Cook, H., Mende, D. R., Letunic, I., Rattei, T., Jensen, L., von Mering, C., and Bork, P. (2018). eggNOG 5.0: a hierarchical, functionally and phylogenetically annotated orthology resource based on 5090 organisms and 2502 viruses. Nucleic Acids Research, 47(D1):gky1085.

Hug, L. A. and Co, R. (2018). It takes a village: Microbial communities thrive through interactions and metabolic handoffs.

Hutchinson, G. E. (1957). Concluding remarks. Cold Spring Harbor Symposia on Quantitative Biology, 22:415–427.

Hyatt, D., Chen, G.-L., Locascio, P. F., Land, M. L., Larimer, F. W., and Hauser, L. J. (2010). Prodigal: prokaryotic gene recognition and translation initiation site identification. BMC Bioinformatics, 11(1):119.

Kanehisa, M., Furumichi, M., Sato, Y., Matsuura, Y., and Ishiguro-Watanabe, M. (2024). KEGG: biological systems database as a model of the real world. Nucleic Acids Res.

Kang, D. D., Li, F., Kirton, E., Thomas, A., Egan, R., An, H., and Wang, Z. (2019). MetaBAT 2: an adaptive binning algorithm for robust and efficient genome reconstruction from metagenome assemblies. PeerJ, 7(e7359):e7359.

Karp, P. D., Paley, S., Krummenacker, M., Kothari, A., Wannemuehler, M. J., and Phillips, G. J. (2022). Pathway Tools Management of Pathway/Genome Data for Microbial Communities. Frontiers in Bioinformatics, 2(April):1–11.

Kochanowski, K., Gerosa, L., Brunner, S. F., Christodoulou, D., Nikolaev, Y. V., and Sauer, U. (2017). Few regulatory metabolites coordinate expression of central metabolic genes in escherichia coli. Molecular Systems Biology, 13(1):903.

Lambert, A., Budinich, M., Mahé, M., Chaffron, S., and Eveillard, D. (2024). Community metabolic modeling of host-microbiota interactions through multi-objective optimization. iScience, 27.

Latorre, C., Betancourt, J. L., Rylander, K. A., and Quade, J. (2002). Vegetation invasions into absolute desert: A 45;th000 yr rodent midden record from the calama–salar de atacama basins, northern chile (lat 22°–24°s). GSA Bulletin, 114(3):349–366.

Lennon, J. T. and Jones, S. E. (2011). Microbial seed banks: the ecological and evolutionary implications of dormancy. Nature reviews microbiology, 9(2):119–130.

Li, D., Liu, C.-M., Luo, R., Sadakane, K., and Lam, T.-W. (2015). MEGAHIT: an ultra-fast single-node solution for large and complex metagenomics assembly via succinct de Bruijn graph. Bioinformatics, 31(10):1674–1676.

Louca, S., Polz, M. F., Mazel, F., Albright, M. B., Huber, J. A., O’Connor, M. I., Ackermann, M., Hahn, A. S., Srivastava, D. S., Crowe, S. A., Doebeli, M., and Parfrey, L. W. (2018). Function and functional redundancy in microbial systems.

Louis, P., Hold, G. L., and Flint, H. J. (2014). The gut microbiota, bacterial metabolites and colorectal cancer. Nat. Rev. Microbiol., 12(10):661–672.

Mandakovic, D., Aguado-Norese, C., García-Jiménez, B., Hodar, C., Maldonado, J. E., Gaete, A., Latorre, M., Wilkinson, M. D., Gutiérrez, R. A., Cavieres, L. A., Medina, J., Cambiazo, V., and Gonzalez, M. (2023). Testing the stress gradient hypothesis in soil bacterial communities associated with vegetation belts in the andean atacama desert. Environmental Microbiome, 18.

Mandakovic, D., Rojas, C., Maldonado, J., Latorre, M., Travisany, D., Delage, E., Bihouée, A., Jean, G., Díaz, F. P., Fernández-Gómez, B., Cabrera, P., Gaete, A., Latorre, C., Gutiérrez, R. A., Maass, A., Cambiazo, V., Navarrete, S. A., Eveillard, D., and González, M. (2018). Structure and co-occurrence patterns in microbial communities under acute environmental stress reveal ecological factors fostering resilience. Scientific Reports, 8.

Mataigne, V., Vannier, N., Vandenkoornhuyse, P., and Hacquard, S. (2021). Microbial systems ecology to understand cross-feeding in microbiomes. Frontiers in Microbiology, 12.

McMurdie, P. and Holmes, S. (2013). phyloseq: An r package for reproducible interactive analysis and graphics of microbiome census data. PLoS ONE, 8:e61217.

Morris, J. J., Lenski, R. E., and Zinser, E. R. (2012). The black queen hypothesis: Evolution of dependencies through adaptive gene loss. mBio, 3.

Muller, E. E., Faust, K., Widder, S., Herold, M., Arbas, S. M., and Wilmes, P. (2018). Using metabolic networks to resolve ecological properties of microbiomes. Current Opinion in Systems Biology, 8:73–80.

Naidoo, Y., Valverde, A., Pierneef, R. E., and Cowan, D. A. (2022). Differences in precipitation regime shape microbial community composition and functional potential in namib desert soils. Microbial Ecology, 83:689–701.

Noor, E., Flamholz, A., Bar-Even, A., Davidi, D., Milo, R., and Liebermeister, W. (2016). The protein cost of metabolic fluxes: prediction from enzymatic rate laws and cost minimization. PLoS computational biology, 12(11):e1005167.

Oksanen, J., Blanchet, F. G., Friendly, M., Kindt, R., Legendre, P., McGlinn, D., Minchin, P. R., O’Hara, R. B., Simpson, G. L., Solymos, P., Stevens, M. H. H., Szoecs, E., and Wagner, H. (2020). vegan: Community Ecology Package. R package version 2.5–7.

Olm, M. R., Brown, C. T., Brooks, B., and Banfield, J. F. (2017). dRep: a tool for fast and accurate genomic comparisons that enables improved genome recovery from metagenomes through de-replication. The ISME Journal, 11(12):2864–2868.

Pacheco, A. R., Moel, M., and Segrè, D. (2019). Costless metabolic secretions as drivers of interspecies interactions in microbial ecosystems. Nature Communications, 10(1):103.

Paine, R. T. (1969). A note on trophic complexity and community stability. The American Naturalist, 103(929):91–93.

Pande, S. and Kost, C. (2017). Bacterial unculturability and the formation of intercellular metabolic networks. Trends in Microbiology, 25:349–361.

Parks, D. H., Imelfort, M., Skennerton, C. T., Hugenholtz, P., and Tyson, G. W. (2015). CheckM: assessing the quality of microbial genomes recovered from isolates, single cells, and metagenomes. Genome Res., 25(7):1043–1055.

Ramos-Tapia, I., Nuñez, R., Salinas, C., Salinas, P., Soto, J., and Paneque, M. (2022). Study of wetland soils of the salar de atacama with different azonal vegetative formations reveals changes in the microbiota associated with hygrophile plant type on the soil surface. Microbiology Spectrum, 10.

Rothman, D. L., Moore, P. B., and Shulman, R. G. (2023). The impact of metabolism on the adaptation of organisms to environmental change. Frontiers in Cell and Developmental Biology, 11:1197226.

Ruscheweyh, H.-J., Milanese, A., Paoli, L., Karcher, N., Clayssen, Q., Keller, M. I., Wirbel, J., Bork, P., Mende, D. R., Zeller, G., and Sunagawa, S. (2022). Cultivation-independent genomes greatly expand taxonomic-profiling capabilities of mOTUs across various environments. Microbiome, 10(1):212.

Régimbeau, A., Budinich, M., Larhlimi, A., Karlusich, J. J. P., Aumont, O., Memery, L., Bowler, C., and Eveillard, D. (2022). Contribution of genome-scale metabolic modelling to niche theory. Ecology Letters, 25:1352–1364.

Saleem, M., Hu, J., and Jousset, A. (2019). More Than the Sum of Its Parts: Microbiome Biodiversity as a Driver of Plant Growth and Soil Health. Annual Review of Ecology, Evolution, and Systematics, 50(1):1–24.

Shade, A., Peter, H., Allison, S. D., Baho, D. L., Berga, M., Bürgmann, H., Huber, D. H., Langenheder, S., Lennon, J. T., Martiny, J. B., Matulich, K. L., Schmidt, T. M., and Handelsman, J. (2012). Fundamentals of microbial community resistance and resilience.

Silverstein, M. R., Bhatnagar, J. M., and Segrè, D. (2024). Metabolic complexity drives divergence in microbial communities. Nat. Ecol. Evol., 8(8):1493–1504.

Sokol, N. W., Slessarev, E., Marschmann, G. L., Nicolas, A., Blazewicz, S. J., Brodie, E. L., Firestone, M. K., Foley, M. M., Hestrin, R., Hungate, B. A., Koch, B. J., Stone, B. W., Sullivan, M. B., Zablocki, O., Trubl, G., McFarlane, K., Stuart, R., Nuccio, E., Weber, P., Jiao, Y., Zavarin, M., Kimbrel, J., Morrison, K., Adhikari, D., Bhattacharaya, A., Nico, P., Tang, J., Didonato, N., Paša-Tolić, L., Greenlon, A., Sieradzki, E. T., Dijkstra, P., Schwartz, E., Sachdeva, R., Banfield, J., and Pett-Ridge, J. (2022). Life and death in the soil microbiome: how ecological processes influence biogeochemistry. Nature Reviews Microbiology, 20:415–430.

Stone, B. W., Li, J., Koch, B. J., Blazewicz, S. J., Dijkstra, P., Hayer, M., Hofmockel, K. S., Liu, X.-J. A., Mau, R. L., Morrissey, E. M., et al. (2021). Nutrients cause consolidation of soil carbon flux to small proportion of bacterial community. Nature communications, 12(1):3381.

Swenson, T. L., Jenkins, S., Bowen, B. P., and Northen, T. R. (2015). Untargeted soil metabolomics methods for analysis of extractable organic matter. Soil Biology and Biochemistry, 80:189–198.

Tay, J. K., Narasimhan, B., and Hastie, T. (2023). Elastic net regularization paths for all generalized linear models. Journal of Statistical Software, 106(1):1–31.

Taş, N., de Jong, A. E., Li, Y., Trubl, G., Xue, Y., and Dove, N. C. (2021). Metagenomic tools in microbial ecology research. Current Opinion in Biotechnology, 67:184–191.

Team, R. C. (2019). R: A Language and Environment for Statistical Computing.

Thommes, M., Wang, T., Zhao, Q., Paschalidis, I. C., and Segrè, D. (2019). Designing metabolic division of labor in microbial communities. mSystems, 4.

Uritskiy, G. V., DiRuggiero, J., and Taylor, J. (2018). MetaWRAP-a flexible pipeline for genome-resolved metagenomic data analysis. Microbiome, 6(1):158.

van der Knaap, J. A. and Verrijzer, C. P. (2016). Undercover: gene control by metabolites and metabolic enzymes. Genes & development, 30(21):2345–2369.

Vikram, S., Guerrero, L. D., Makhalanyane, T. P., Le, P. T., Seely, M., and Cowan, D. A. (2016). Metagenomic analysis provides insights into functional capacity in a hyperarid desert soil niche community. Environmental microbiology, 18:1875–1888.

Vásquez-Dean, J., Maza, F., Morel, I., Pulgar, R., and González, M. (2020). Microbial communities from arid environments on a global scale. A systematic review. Biological Research, 53(1):29.

Wang, H., Bu, L., Tian, J., Hu, Y., Song, F., Chen, C., Zhang, Y., and Wei, G. (2021). Particular microbial clades rather than total microbial diversity best predict the vertical profile variation in soil multifunctionality in desert ecosystems. Land Degradation and Development, 32:2157–2168.

Wang, X., Xia, K., Yang, X., and Tang, C. (2019). Growth strategy of microbes on mixed carbon sources. Nature Communications, 10.

Wang, X.-W., Sun, Z., Jia, H., Michel-Mata, S., Angulo, M. T., Dai, L., He, X., Weiss, S. T., and Liu, Y.-Y. (2024). Identifying keystone species in microbial communities using deep learning. Nat. Ecol. Evol., 8(1):22–31.

Wang, Y.-P. and Lei, Q.-Y. (2018). Metabolite sensing and signaling in cell metabolism. Signal transduction and targeted therapy, 3(1):30.

Wickham, H. (2016). ggplot2: Elegant Graphics for Data Analysis.

Wu, Y.-W., Simmons, B. A., and Singer, S. W. (2015). MaxBin 2.0: an automated binning algorithm to recover genomes from multiple metagenomic datasets. Bioinformatics, 32(4):605–607.

Xun, W., Liu, Y., Li, W., Ren, Y., Xiong, W., Xu, Z., Zhang, N., Miao, Y., Shen, Q., and Zhang, R. (2021). Specialized metabolic functions of keystone taxa sustain soil microbiome stability. Microbiome, 9.

Zhou, Z., Tran, P. Q., Breister, A. M., Liu, Y., Kieft, K., Cowley, E. S., Karaoz, U., and Anantharaman, K. (2022). Metabolic: high-throughput profiling of microbial genomes for functional traits, metabolism, biogeochemistry, and community-scale functional networks. Microbiome, 10.

Ziesack, M., Gibson, T., Oliver, J. K. W., Shumaker, A. M., Hsu, B. B., Riglar, D. T., Giessen, T. W., DiBenedetto, N. V., Bry, L., Way, J. C., Silver, P. A., and Gerber, G. K. (2019). Engineered interspecies amino acid cross-feeding increases population evenness in a synthetic bacterial consortium. mSystems, 4(4).

Øyvind Hammer, Harper, D. A., and Ryan, P. D. (2001). Past: Paleontological statistics software package for education and data analysis. Palaeontologia Electronica, 4.

