## Supplementary file 1 - Figures and Text for "Modeling the emergent metabolic potential of soil microbiomes in Atacama landscapes"

#### Supplementary Material

- **Excel file:**
  - **Table S1:** Nutritional and physicochemical metadata.
  - **Table S2:** Sequencing information of metagenomes.
  - **Table S3:** Raw, relative, and hellinger-transformed abundances of operational taxonomic units.
  - **Table S4:** Raw, relative, and hellinger-transformed abundances of taxonomic ranks from phyla to genra calculated upon Table S3. SIMPER analyses were calculated upon the hellinger-transformed data.
  - **Table S5:** Raw, relative, and hellinger-transformed abundances of functional annotations of genes. SIMPER analyses were calculated upon the hellinger-transformed data.
  - **Table S6:** Genomic information of assembled MAGs.
  - **Table S7:** Merged reports of unique reactions predicted per site by dataset.
  - **Table S8:** Nutritional seeds provided as input in the network expansion step.
  - **Table S9:** Binary matrix for producible metabolites per simulated condition (site x seed) and dataset.
  - **Table S10:** Metabolite groups with non-zero coefficients and, thus, fitted as relevant for at least one environmental variables when targeted with the elastic net.
  - **Table S11:** Producible targets due to individual or cooperative efforts by site.
- **Figures S1 to S5**
- **Table S12:** Reactions in metabolic networks
- **Extended Results:**
  - Background of the TLT
    - Suppl. Text S1: Environmental description
    - Suppl. Text S2: Taxonomic description
    - Suppl. Text S3: Functional description
    - Suppl. Text S4: Crossing taxonomic and functional insights
  - Metabolism of the TLT
    - Suppl. Text S5: Nitrogen cycling at the lagoon (EV1)
    - Suppl. Text S6: Carbon cycling at the puna (EV2)
    - Suppl. Text S7: Taxonomy of key species in minimal communities
- **Supplementary bibliography**

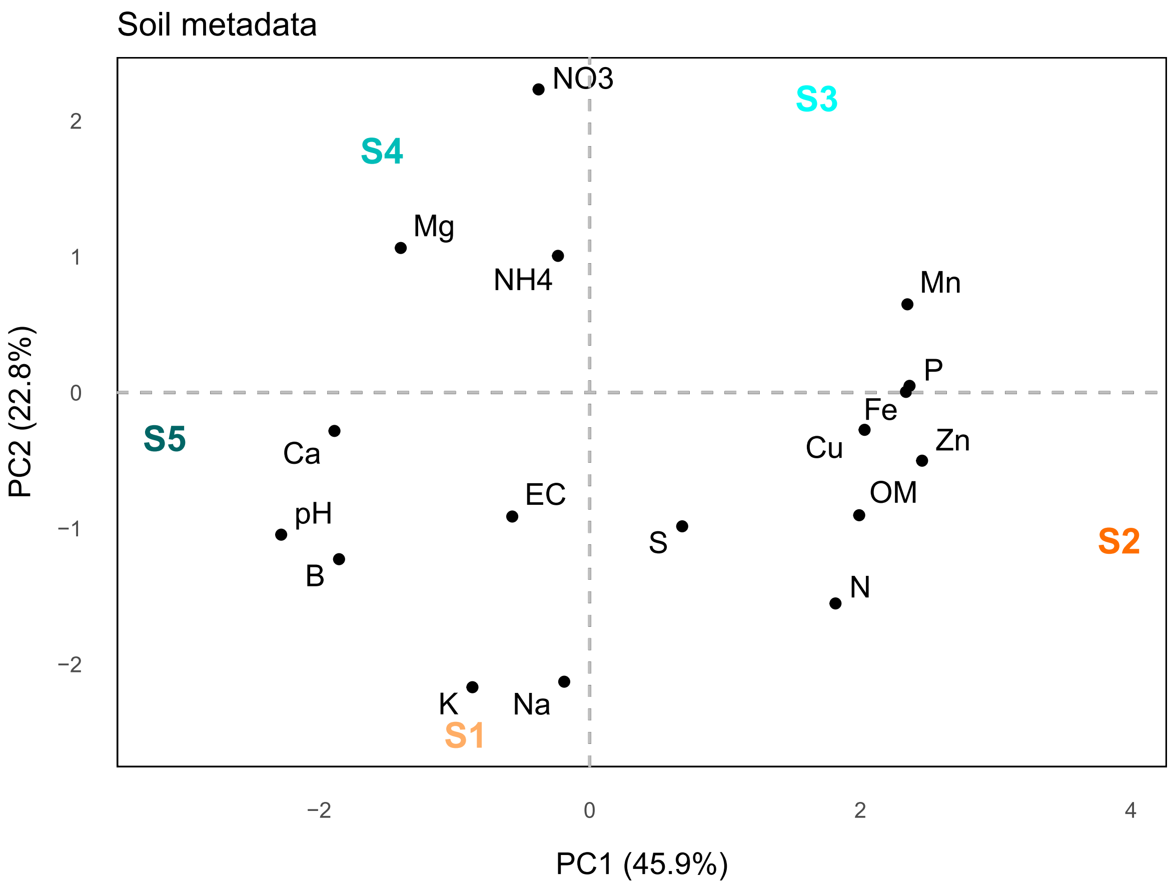


Figure S1: PCA soil metadata excluding S6.

### 1 Background of the TLT

#### Supp. Text S1: Environmental description of the TLT (related to Fig. S1).

In this study, sites S1 and S2 survey prepuna and puna ecosystems at +2800 and +3800 meters above sea level (m.a.s.l.), respectively. While S1 was highly saline, with a pH of 8.1, and mainly driven by high contents of potassium (K) and sodium (Na), S2 exhibited a pH of 6.1 and high measurements for multiple environmental variables including zinc (Zn), copper (Cu), iron (Fe), sulfur (S), nitrogen (N), and organic matter (OM, Fig. S1). In particular, measurements of environmental N (60±30 mg/kg) and OM (1.02±0.03 %) at S2 were significantly higher than in any other site. Sites S3, S4, and S5 span steppe ecosystems (+4300 m.a.s.l.) with pH measurements ranging from 5.8 to 8.5 (Suppl. Table 1). Site S3, under the nutritional influence of manganese (Mn) and nitrate ($NO_3$), is the most acidic soil; S4 showed a neutral pH value (7.5) and high contents of magnesium (Mg) and ammonium ($NH_4$); and S5 was basic and characterized by calcium (Ca) and boron (B, Fig. S1). Lastly, site S6 was collected from the Lejía lagoon's shore, constituting an ecosystem of its own, hypersaline, that behaved as an outlier for phosphorus (P, 59±2 mg/kg), K (770±30 mg/kg), Mg (1362±81 mg/kg), S (488.5±12.5 mg/kg), Na (982±40 mg/kg), B (98.3±2.7 mg/kg), Fe (169.5±3.5 mg/kg), and electric conductivity (EC, 1.94±0.15 mS/cm, Fig. 4B) explaining its overall dissimilarity from the rest of sites (Fig. 1A).

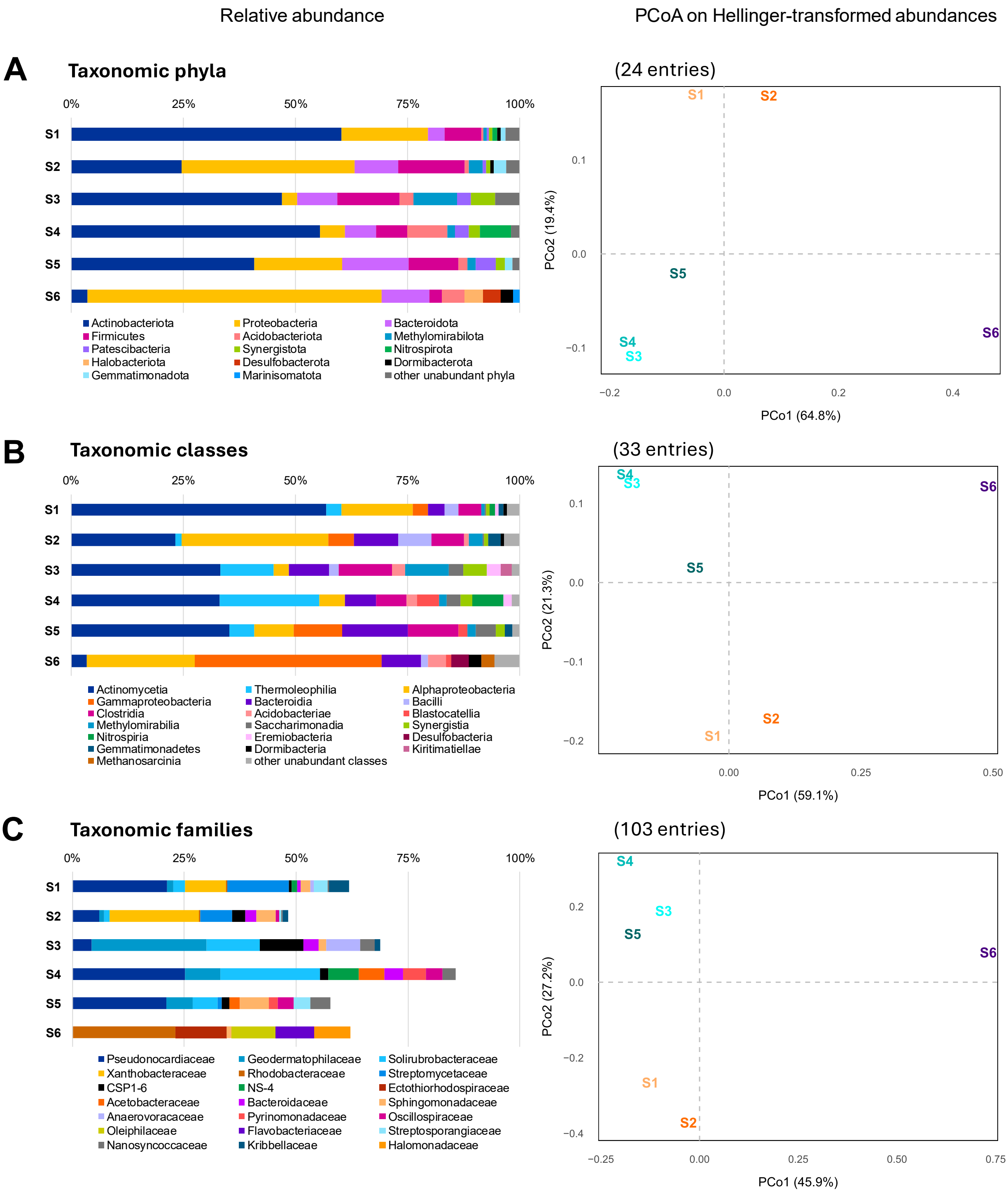


Figure S2: **Taxonomic profiling and ordination analyses show clear and consistent separation of ecosystems across ranks. (left)** Relative abundance of taxonomic phyla (A), classes (B), and families (C). Taxa with abundance < 1% and < 2% in all sites were merged into 'other unabundant' phyla and classes, respectively. Taxonomic families contributing up to 40% of the cumulative overall dissimilarity according to a Similarity percentage analysis (SIMPER) are shown. Ranked importance decreases from left to right. **(right)** Principal coordinates analysis (PCoA) conducted upon the Hellinger-transformed abundances.

#### Supp. Text S2: Taxonomic description of the TLT (related to Fig. S2).

Taxonomic profiling of the studied prokaryotic communities revealed different microbial composition across sites even at the levels of phylum and class (Figs. S2A,B). Overall and consistent with what it is expected in deserts(Feng et al., 2020; Naidoo et al., 2022), soils from the TLT are dominated by bacteria from Actinobacteriota and Proteobacteria phyla, with average relative abundances of 38.6%±21.2% and 25.3%±23.4%, respectively, followed by Firmicutes (9.6%±4.6%), Bacteroidota (9.1%±3.7%), and Acidobacteriota (3.5%±3.2%). However, examination down to the rank of orders revealed little overlap of taxa between the surveyed ecosystems (Fig. 2B). For instance, while most common Actinobacteriota across the TLT belong to Mycobacteriales (that include Pseudonocardiales, according to GTDB's taxonomic opinion), the prepuna (S1) and puna (S2) are enriched in Streptomycetales and Actinomycetales, sites from the steppe (S3 to S5) are so in Solirubrobacterales, and the sediment microbiome from S6 barely harbors members from this phylum. Another example is the distribution of Rhizobiales, restricted to S1 and S2, Rhodobacterales that dominate in S6 and Pseudomonodales and Ectothiorhodospirales that were exclusively found in that sample, while all these taxa from the Proteobacteria phylum were absent in the steppe microbiomes. Our results are supported by other studies conducted on desert soils that pointed Solirubrobacterales and Actinomycetales, important decomposers of organic matter, as highly abundant (McHugh et al., 2017; Crits-Christoph et al., 2013; Hung et al., 2022) and members from Streptomycetales as predictors of soil chemistry at the puna belt of the TLT (Mandakovic et al., 2023) while they further underpin the differing importance of such taxa in the microbiome structure of the sampled ecosystems. In particular, Solirubrobacterales have been found in steppes of Mongolia and China (Qin et al., 2021; Song et al., 2023) and associated to heavily disturbed soils (Shange et al., 2012). On the other hand, members from Rhizobiales have been described as nitrogen suppliers commonly occurring in semiarid ecosystems, even in bare soils, and found to comprise most Alphaproteobacteria in the Mojave Desert (McHugh et al., 2017). This suggests that the high abundance observed in the prepuna and puna, that exhibit high levels of environmental N, reflect an indispensable contribution of Rhizobiales to the whole communities they belong to. Additionally, Rhodobacterales have been associated to carbon-fixing pathways in microbial mats from hypersaline lakes in Atacama, specifically to pathways of sulfur oxidation and anoxygenic photosynthesis (Kurth et al., 2021), allowing to speculate that members from this order provide fundamental strategies for microbiome functioning in the demanding conditions that soils from the Lejía lagoon are exposed to.

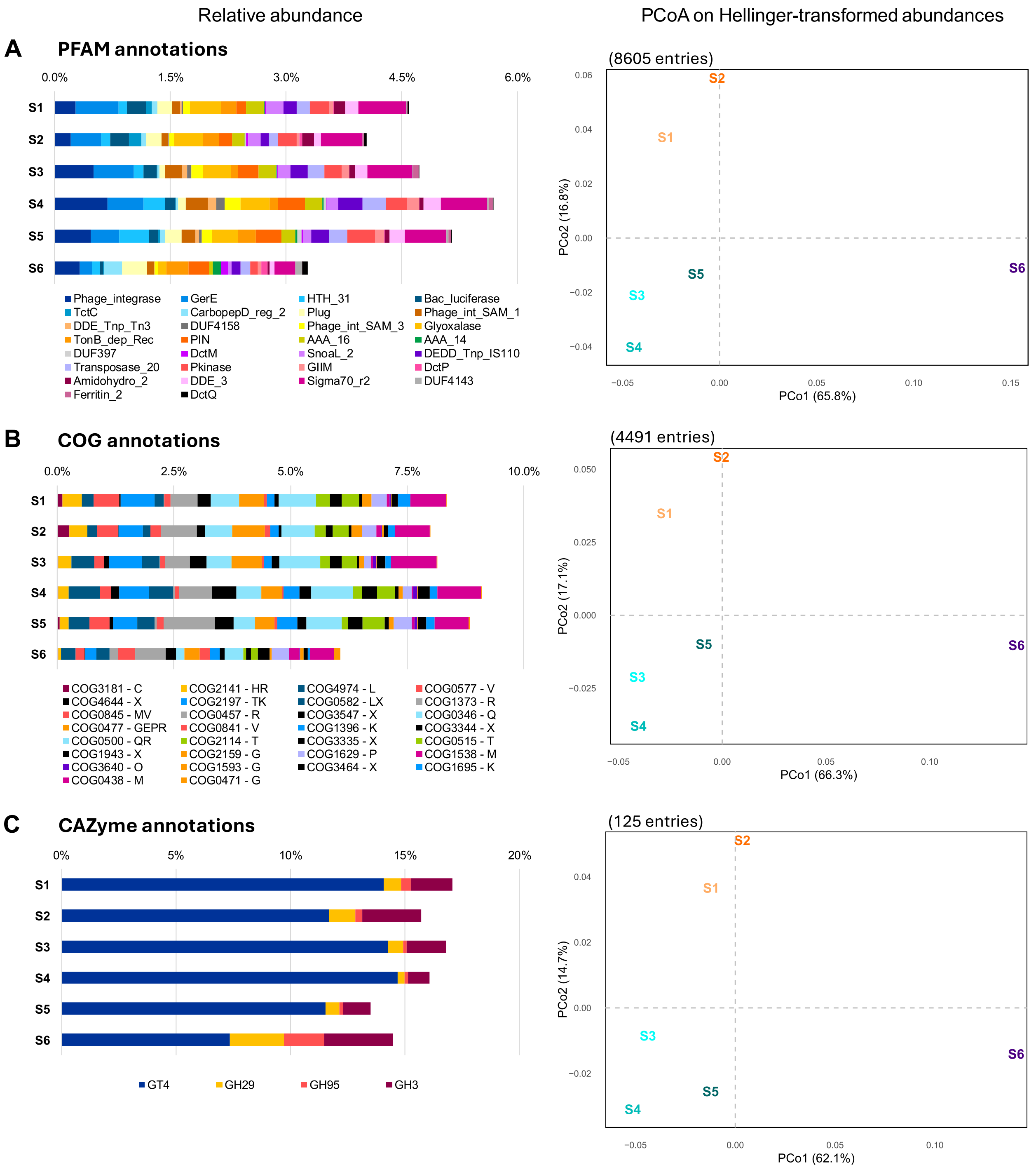


Figure S3: **Functional profiling upon PFAM and COG annotations evidence overall consistency while ordination analyses capture the separation of ecosystems observed in Figs. 2 and S1. (left)** Relative abundance of most dissimilar functions between sites following the PFAM (A) and COG (B and C) annotations. The top 30 ranked functions contributing to the cumulative overall dissimilarity according to a Similarity percentage analysis (SIMPER) are shown in (A) and (B). Importance decreases from left to right. Capital letters after hyphens in (B) indicate the COG category which each annotation belongs to. 'Other categories' in (C) comprise functions annotated as W, F, Z, B, A, and Y. **(right)** Principal coordinates analysis (PCoA) conducted upon the Hellinger-transformed abundances.

#### Supp. Text S3: Functional description of the TLT (related to Fig. S3).

Our results show that the prepuna (S1) and puna (S2) microbiomes are enriched in annotations for degradation of aromatic compounds (ko01220) such as benzoate (ko00362), aminobenzoate (ko00627), and chlorobenzoate (ko00361), complementing previous evidence of enriched pathways for 3-phenylpropanoate and aerobic toluene degradation in soils from the Atacama Desert (Ramos-Tapia et al., 2022). In addition, S6 was characterized by KEGG pathways related to flagellar assembly (ko02040), chemotaxis (ko02030), and two-component systems (ko02020). We speculate that the enrichment of these functions in the microbiome collected from the Lejía lagoon's shore is due to the need of motility in hypersaline conditions where osmotic stress constantly threatens cellular integrity. Chemotaxis allows microorganisms to detect chemical stimuli and move accordingly through the display of membrane receptors, typically two-component systems, that sense the environment and initiate the signal transduction. Ultimately, this affects flagellar rotation and induces the movement towards or away from the chemical gradient, allowing microorganisms to locate nutrients and to avoid harmful substances (Normand et al., 2015). This conjecture sets motility as a crucial trait in this ecosystem, considering the ongoing evaporation of the lagoon and sporadic rewetting events.

Carbohydrate-active enzymes (CAZymes) inform on carbon metabolism across the samples. The broad family of glycosyl transferases GT4 accounted, in average, for 13.3% of relative abundance calculated upon CAZyme annotations in S1 to S5, while reduced to almost half in S6 (7.4%). On the other hand, this sample was enriched in glycosyl hydrolases from the families GH29 and GH95 —that include galactosidases and fucosidases— and that together with the broad GH3 family account for 7.1% of relative abundance whereas sites S1 to S5 averaged in total a content of 2.6% for these annotations (Fig. S3C). These results suggest that galactose and fucose likely comprise relevant carbon sources in the nutrient-poor shore of the Lejía lagoon and that the enzymes responsible for the breakdown of substrates containing them must handle high salinity. Although glycosyl hydrolases have been reported in extreme environments, knowledge about their functions in saline deserts is limited (Ashcroft and Munoz-Munoz, 2024). Finally, while galactosidases and fucosidases have been associated to acidophiles (Ashcroft and Munoz-Munoz, 2024) and forest soils (Houfani et al., 2019), our results indicate that at they might help degrade extracellular polymeric substances synthesized by the microbial community from S6 (Lennon and Jones, 2011). This context sets the TLT as a promising opportunity to unveil how microbes obtain energy in extreme conditions.

To address non-metabolic features potentially characterizing the steppe microbiomes, we deepened into PFAM (Fig. S3A) and COG (Fig. S3B) annotations and found overall consistency between both nomenclatures. Main results show an enrichment in steppe samples, particularly in S4, of mobile genetic elements including transposases (e.g., COG4644, COG3547, COG3335, COG3464, Transposase_20, DDE_Tnp_Tn3, DEDD_Tnp_IS110, DUF4158), phage integrases (e.g., COG0582, Phage_int_SAM_1, Phage_int_SAM_3), and other related domains like reverse transcriptases (e.g., COG3344, GIIM), site-specific recombinases (e.g., COG4974), and tetratricopeptide repeats (e.g. COG0457, Figs. S2 A and S2 B). At the coarse resolution of COG categories, these findings are reflected by the enrichment in the steppe samples of the L category that is defined by functions related to replication, recombination, and DNA repair (data not shown). These results agree with a previous report showing the enrichment of mobile genetic elements and their co-occurrence with the L COG category in hyperarid sites of the Atacama Desert (Sáenz et al., 2019) and suggest that the environmental stressors that shape steppes from the TLT constitute extremely demanding conditions to sustain life. Indeed, Sáenz and colleagues hypothesized that bacterial communities with low taxonomic richness, like the observed in S4, resort to extensive gene exchange mechanisms to increase genomic diversity, where a potential for gene duplication could represent an adaptative advantage to cope, for example, with the increased UV-induced damage expected in the studied Atacama high-lands (+4,000 m.a.s.l.) Sáenz et al. (2019). Finally, this examination revealed that the most dissimilar COG annotation across the complete dataset corresponded to tripartite-type tricarboxylate transporters (COG3181) which was enriched in S1 and S2, particularly in the puna sample, result supported by the PFAM-based approach (TctC). This observation is consistent with reports of an enriched pathway for reductive tricarboxilic acid cycle I in soils from the high puna in the Atacama Desert (Ramos-Tapia et al., 2022). Despite of being found in steppe soils, another study evidenced the enrichment of genes annotated as COG3181 which depicts the relevance of these carbon transporters across arid and semiarid soils (Song et al., 2023). Altogether, the latter supports the result obtained through the KEGG-based approach, further pointing out the puna microbiome from S2 as a community of enhanced capabilities for the uptake and catabolism of organic carbon sources.

#### Supp. Text S4: Crossing taxonomic and functional insights (related to Figs. S2 and S3).

Given the conserved distribution of samples across taxonomies and functionalities (Figs. 2B and 2C), and leaning on other studies conducted in deserts that showed strong correlations between phylogenetic and functional diversities (Naidoo et al., 2022; Fierer et al., 2012), we propose that dissimilar abundances of taxonomic families might be responsible for the genes highlighted as distinct between sites in the previous section. So, to couple microbial community composition with most dissimilar functions, through a SIMPER analysis performed as above, we ranked most dissimilar taxonomic families (Fig. S2C) and found sharp differences between sites. Specifically, Xanthobacteraceae was exclusively detected in the prepuna and puna, enabling to hypothesize that members from this taxon may be the ones accounting for the enhanced potential for the degradation of aromatic compounds in S2 and, to a lesser extent, in S1. In fact, this family has already been inferred to handle p-nitrophenol degradation (Cao et al., 2022) and to encode for a flavin monooxygenase (4-Hydroxyphenylacetate 3-hydroxylase) capable to catalize aromatic ring hydrolization (Chen et al., 2021). Furthermore, recently, a strain from Xanthobacteraceae, isolated from the rhizosphere of a plant used in phytorremediation of policyclic aromatic compounds, was proven to degrade pyrene at pH 5 to 7 (Ge et al., 2024) which sets this function as prone to emerge in the sampled puna microbiome that exhibited a pH value of 6.1. Related to steppes of the TLT, besides the family Solirubrobacteraceae that comprissed all Solirubrobacterales in S3 to S5, our results underpin the role of Geodermatophilaceae in the increased potential for DNA repair and recombination observed in those samples. This family has been associated with transposable elements in both genomic (Sghaier et al., 2016) and metagenomic (Salam et al., 2017) studies. Also, a systematic study on the pseudogenes of the suborder Frankineae, classified as Mycobacteriales according to the taxonomic opinion from GTDB, revealed that Geodermatophilaceae had a significant portion of these disabled copies of functional genes compared to other families from the same taxon which indicates an evolutionary history of failed attempts of horizontal gene transfer (Sur et al., 2013). Finally, the rare taxonomic and functional profiles from S6 deliver a more complex outline to unveil the relationship between both layers of information as they provide several taxonomic families as possible candidates for dissimilar functions and we abstain of further speculations. Nonetheless, our analyses advocate for conceiving particular microbial populations, not necessarily abundant, as major role players in community growth, functioning, and ecosystem interactions (Wang et al., 2021).

### 2 Metabolism of the TLT

| Sites | Metagenomes (%) | Union of MAGs (%)* |
|---|---|---|
| S1 | 5,884 (83.0) | 2,820 (75.2) |
| S2 | 6,261 (83.0) | 4,233 (74.7) |
| S3 | 5,735 (83.1) | 3,594 (75.2) |
| S4 | 5,700 (82.5) | 3,579 (75.0) |
| S5 | 5,803 (82.7) | 3,935 (76.0) |
| S6 | 5,587 (82.9) | 4,258 (74.0) |
| Average | 5,828.3 (82.8) | 3736.5 (75.0) |
| St. dev. | 234.3 (0.2) | 537.2 (0.7) |

**Table S12.** Summary of reactions in input metabolic networks. Values in brackets indicate percentage of gene-related reactions. *Percentage of gene-related reaction for MAGs was calculated upon average of individual genome-resolved metabolic networks by site. Details on individual reports are detailed in Tables S6 and S7.

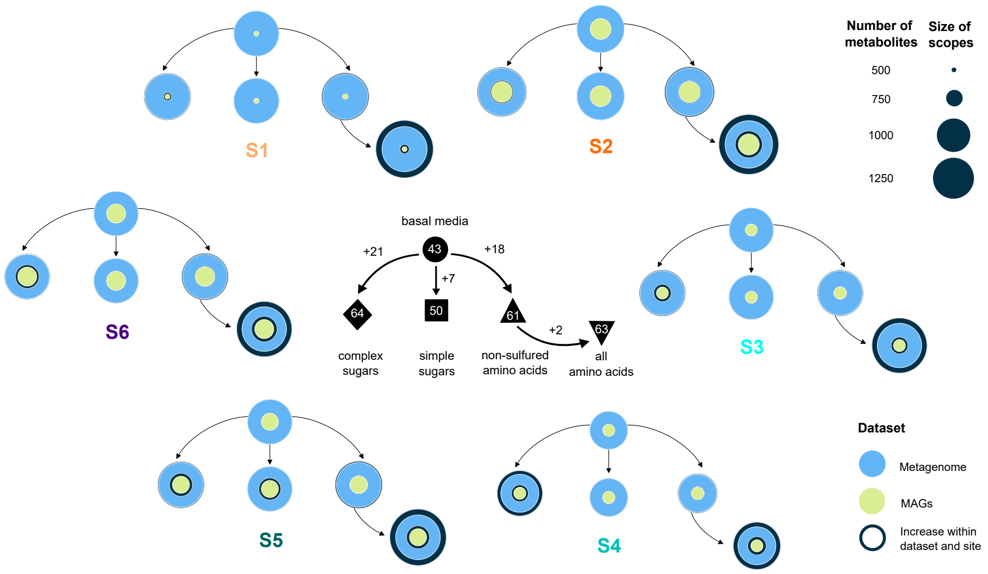


Figure S4: **Quantitative description of scopes.** Five seeds simulating environmental availability of inorganic carbon (basal media), organic carbon (simple and complex sugars), organic nitrogen (non-sulfured amino acids), and organic sulfur (all amino acids) were provided as input for the expansion of metabolic networks. Numbers inside shapes at the central flowchart represent the number of compounds in each user-provided seed and numbers next to arrows show the number of compounds added on top of the previous seed. The basal media is constituted by coenzymes (n=18), metal ions (n=6), and inorganic chemicals (n=19) that include oxygen, water, carbon dioxide, bicarbonate, ammonium, nitrate, nitrite, sulfate, sulfite, and thiosulfate. The output obtained after the network expansion with Metage2Metabo are lists of metabolites predicted to be producible (scopes) which are quantitatively represented by the area of colored spheres for each site, labeled from S1 to S6 in clockwise direction. Outlines in charcoal blue depict the increase in sizes of scopes for each seed compared to the *basal media* by dataset within site, outer: metagenomes, inner: MAGs.

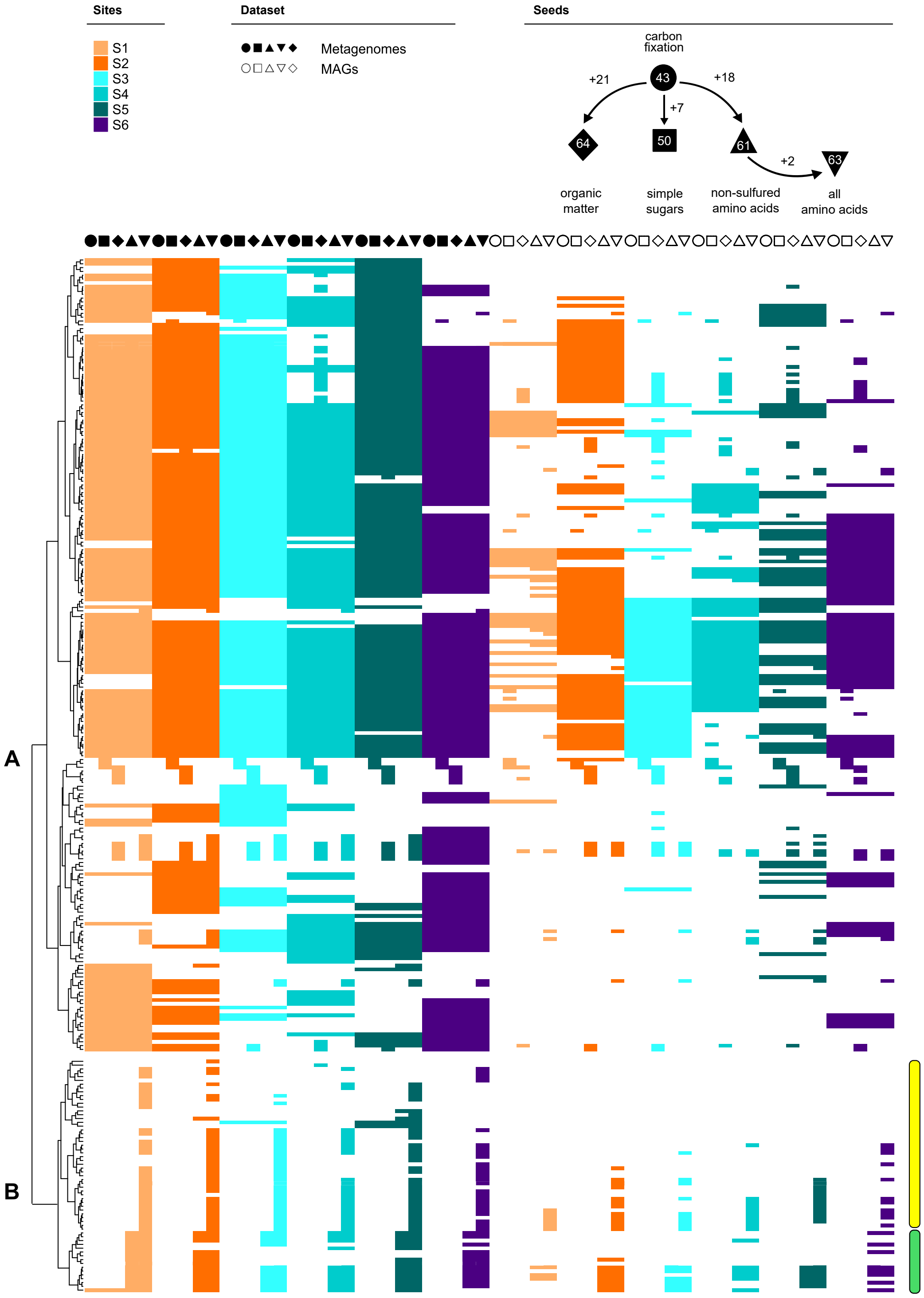


Figure S5: **Overview of the metabolic potential**. Metabolite groups (n=269) for the union of all reachable metabolites across sites, seeds, and datasets (n=1517). (A) Metabolite groups that hold distinctions between sites with minor impact of nutritional seeds within sites (n=211). The upper left quadrant reflects the redundant gene reservoir that metagenomes shelter across sites. The upper right quadrant reflects the patched functional overlaps of the MAG-scale metabolic networks that characterize their site-driven behavior. (B) Metabolite groups related to the effect of amino acids (n=58). Color bars at the right indicate metabolites mainly reachable due to the effect of non-sulfured amino acids (green) and of cysteine and methionine (yellow) highlighting the role of organic nitrogen and organic sulfur, respectively, when environmentally available. See last column of Table S9 for details on individual metabolites.

#### Supp. Text S5: The Lejía lagoon is under several environmental pressures associated to osmotic stress and nitrogenated pathways (related to Fig. 4).

The first group of environmental variables (EV1) was defined by K, Na, Fe, P, Mn, Mg, S, B, Ca and electric conductivity, all for which S6 was the site with highest measurements, implying that abundance of these elements are likely to constitute major abiotic stressors for microorganisms inhabiting the rare ecosystem found in the shore of the surveyed brine lagoon. The elastic net model positively associated these ten variables to five metabolite groups representing 23 metabolites that have previously been linked to nitrogenated pathways and osmotic stress. Specifically, isovalerate and isobutyrate are involved in L-leucine II and IV and L-valine III degradation while 2-methylbutanoate is involved in L-isoleucine V biosynthesis, all wrapped in X3_P009. Also, 4-guanidinobutanoate (X1_P097) is involved in L-arginine degradation and aceneuramate (X17_P000) is involved in N-acetylneuraminate and N-acetylmannosamine degradation. On the other hand, the chemical structures of some of these nitrogenated compounds classifies them as short chain fatty acids (2-methylbutanoate and isovalerate) and glycans (e.g. N-Actyl-D-glucosamine, N-acetyl-$\beta$-D-glucosaminyl-(1$\rightarrow$4)-D-glucosamine, and chitobiose in X17_P000, only producible in MetaG-GEMs). Furthermore, $\beta$-glutamate (X1_P040) is highly associated with both potassium and sodium, highlighting its relevance as an osmoprotective agent in arid soils (Dinnbier et al., 1988).

#### Supp. Text S6: Microbes from the puna are specialized in carbon-cycling, possibly due to high contents of zinc, nitrogen, and organic matter (related to Fig. 4).

The second group of environmental variables (EV2) was defined by OM, N, and Zn for which S2 registered the highest in situ measurements and was an outlier for the first two. Two metabolite groups were related to all drivers that this group gave off and consisted of D-ribulose (X1_P103), a sugar formed in the pentose phosphate pathway that serves as a precursor for several bioactive compounds including nucleotides implying a relevance on DNA repair (Leung et al., 2020) and fifteen metabolites in X15_P000 including chromopyrrolate and 2-imino-3-(indol-3-yl)propanoate, protodeoxyviolaceinate and prodeoxyviolacein, and naringenin chalcone, involved in staurosporine, violacein, and flavonoid biosynthesis, respectively, antioxidants that could protect these microbial communities against oxidative stress (Georgiou et al., 2015) and UV radiation (Knaggs, 2000). On the other hand, (2Z,4Z)-2-hydroxy-5-carboxymuconate-6-semialdehyde (X1_P071), involved in the degradation of 4-amino-3-hydroxybenzoate and protocatechuate (III), associated to OM and N but not to Zn. Additionally, constituted by (+)-epi-isozizaene, a terpenoid involved in albaflavenone biosynthesis, guanidinium, a precursor of arginine and urea biosynthesis, and a quinone, two amino acids, and a dipeptide; X6_P002 was associated to environmental N only. These results indicate that microbes from the surveyed puna ecosystem harbor metabolic capabilities for the catabolism of aromatic compounds and other organic molecules, consistent with our results obtained from the functional profiling that show enhanced metabolic potential for carbon cycling in this site.

#### Supp. Text S7: Taxonomy of key species in minimal communities (related to Fig. 5).

We analysed the taxonomy of MAGs occurring in minimal communities (Fig. 5). MAGs from Actinomycetota (Actinobacteriota) constitute key species in all the sampled sites. With at least one MAG with relative abundance > 1% in all minimal communities from S1 to S5, the vast majority were essential symbionts implying that the functional contributions of this phylum to the cooperative efforts for baring the environmental pressures of the TLT are irreplaceable. The only exception was observed in S6 where MAGs from Actinomycetota were unabundant and classified as alternative symbionts; however, the possible replacement was another MAG from the same taxon. Next, we found that MAGs from Acidobacteriota were essential in S3 to S5, and that they can serve as replacement for bacteria classified as Verrucomicrobiota (S2 and S4), Bacteroidota (S4), and Thermoproteota (S5). The latter underscores the documented metabolic diversity of Acidobacteriota (Sikorski et al., 2022; Waschulin et al., 2022; Crits-Christoph et al., 2022) which seems to enable members from this phylum to successfully adapt to the stressful steppe deserts of the TLT and to constitute a reserve of functions, playing a major role in sustaining the stability of such soil communities subjected to permanent abiotic perturbations (Díaz et al., 2016). Additionally, we observed that MAGs from Dormibacterota were key species in the slightly acidic sites of S2 and S3 which exhibited pH values of 6.1 and 5.8, respectively. Finally, some phyla were essential in minimal communities of an exclusive site; for example, Myxococcota was key only in the puna ecosystem (S2) while Desulfobacterota and Bacillota (Firmicutes) were essential symbionts only in the Lejía lagoon's shore (S6).

### References


Ashcroft, E. and Munoz-Munoz, J. (2024). A review of the principles and biotechnological applications of glycoside hydrolases from extreme environments. *International Journal of Biological Macromolecules*, 259.

Cao, L., Zhu, G., Tao, J., and Zhang, Y. (2022). Iron carriers promote biofilm formation and p-nitrophenol degradation. *Chemosphere*, 293.

Chen, R., Miao, Y., Liu, Y., Zhang, L., Zhong, M., Adams, J. M., Dong, Y., and Mahendra, S. (2021). Identification of novel 1,4-dioxane degraders and related genes from activated sludge by taxonomic and functional gene sequence analysis. *Journal of Hazardous Materials*, 412.

Crits-Christoph, A., Diamond, S., Al-Shayeb, B., Valentin-Alvarado, L., and Banfield, J. F. (2022). A widely distributed genus of soil acidobacteria genomically enriched in biosynthetic gene clusters. *ISME Communications*, 2.

Crits-Christoph, A., Robinson, C. K., Barnum, T., Fricke, W. F., Davila, A. F., Jedynak, B., McKay, C. P., and DiRuggiero, J. (2013). Colonization patterns of soil microbial communities in the Atacama Desert. *Microbiome*, 1(1):28.

Dinnbier, U., Limpinsel, E., Schmid, R., and Bakker, E. P. (1988). Transient accumulation of potassium glutamate and its replacement by trehalose during adaptation of growing cells of escherichia coli k-12 to elevated sodium chloride concentrations. *Archives of Microbiology*, 150:348–357.

Díaz, F. P., Frugone, M., Gutiérrez, R. A., and Latorre, C. (2016). Nitrogen cycling in an extreme hyperarid environment inferred from $\delta^{15}$N analyses of plants, soils and herbivore diet. *Scientific Reports*, 6(22226).

Feng, W., Zhang, Y., Yan, R., Lai, Z., Qin, S., Sun, Y., She, W., and Liu, Z. (2020). Dominant soil bacteria and their ecological attributes across the deserts in northern china. *European Journal of Soil Science*, 71:524–535.

Fierer, N., Leff, J. W., Adams, B. J., Nielsen, U. N., Bates, S. T., Lauber, C. L., Owens, S., Gilbert, J. A., Wall, D. H., and Caporaso, J. G. (2012). Cross-biome metagenomic analyses of soil microbial communities and their functional attributes. *Proceedings of the National Academy of Sciences of the United States of America*, 109:21390–21395.

Ge, H., Liu, X., Lu, D., Yang, Z., and Li, H. (2024). Degradation of pyrene by xanthobacteraceae bacterium strain s3 isolated from the rhizosphere sediment of vallisneria natans: active conditions, metabolite identification, and proposed pathways. *Environmental Science and Pollution Research.*

Georgiou, C. D., Sun, H. J., McKay, C. P., Grintzalis, K., Papapostolou, I., Zisimopoulos, D., Panagiotidis, K., Zhang, G., Koutsopoulou, E., Christidis, G. E., et al. (2015). Evidence for photochemical production of reactive oxygen species in desert soils. *Nature Communications*, 6(1):7100.

Houfani, A. A., Větrovský, T., Navarrete, O. U., Štursová, M., Tláskal, V., Beiko, R. G., Boucherba, N., Baldrian, P., Benallaoua, S., and Jorquera, M. A. (2019). Cellulase-hemicellulase activities and bacterial community composition of different soils from algerian ecosystems. *Microbial Ecology*, 77:713–725.

Hung, C. M., Chen, C. W., Huang, C. P., and Dong, C. D. (2022). Degradation of 4-nonylphenol in marine sediments using calcium peroxide activated by water hyacinth (eichhornia crassipes)-derived biochar. *Environmental Research*, 211.

Knaggs, A. R. (2000). The biosynthesis of shikimate metabolites. *Natural Product Reports*, 17:269–292.

Kurth, D., Elias, D., Rasuk, M. C., Contreras, M., and Farias, M. E. (2021). Carbon fixation and rhodopsin systems in microbial mats from hypersaline lakes brava and tebenquiche, salar de atacama, chile. *PLoS ONE*, 16.

Lennon, J. T. and Jones, S. E. (2011). Microbial seed banks: the ecological and evolutionary implications of dormancy. *Nature reviews microbiology*, 9(2):119–130.

Leung, P. M., Bay, S. K., Meier, D. V., Chiri, E., Cowan, D. A., Gillor, O., Woebken, D., and Greening, C. (2020). Energetic basis of microbial growth and persistence in desert ecosystems. *Msystems*, 5(2):10–1128.

Mandakovic, D., Aguado-Norese, C., García-Jiménez, B., Hodar, C., Maldonado, J. E., Gaete, A., Latorre, M., Wilkinson, M. D., Gutiérrez, R. A., Cavieres, L. A., Medina, J., Cambiazo, V., and Gonzalez, M. (2023). Testing the stress gradient hypothesis in soil bacterial communities associated with vegetation belts in the andean atacama desert. *Environmental Microbiome*, 18.

McHugh, T. A., Compson, Z., van Gestel, N., Hayer, M., Ballard, L., Haverty, M., Hines, J., Irvine, N., Krassner, D., Lyons, T., Musta, E. J., Schiff, M., Zint, P., and Schwartz, E. (2017). Climate controls prokaryotic community composition in desert soils of the southwestern united states. *FEMS Microbiology Ecology*, 93.

Naidoo, Y., Valverde, A., Pierneef, R. E., and Cowan, D. A. (2022). Differences in precipitation regime shape microbial community composition and functional potential in namib desert soils. *Microbial Ecology*, 83:689–701.

Normand, P., Caumette, P., Goulas, P., Pujic, P., and Wisniewski-Dyé, F. (2015). *Adaptations of Prokaryotes to Their Biotopes and to Physicochemical Conditions in Natural or Anthropized Environments*, pages 293–351. Springer Netherlands, Dordrecht.

Qin, J., Li, M., Zhang, H., Liu, H., Zhao, J., and Yang, D. (2021). Nitrogen deposition reduces the diversity and abundance of cbbl gene-containing co2-fixing microorganisms in the soil of the stipa baicalensis steppe. *Frontiers in Microbiology*, 12.

Ramos-Tapia, I., Nuñez, R., Salinas, C., Salinas, P., Soto, J., and Paneque, M. (2022). Study of wetland soils of the salar de atacama with different azonal vegetative formations reveals changes in the microbiota associated with hygrophile plant type on the soil surface. *Microbiology Spectrum*, 10.

Salam, L. B., Obayori, S. O., Nwaokorie, F. O., Suleiman, A., and Mustapha, R. (2017). Metagenomic insights into effects of spent engine oil perturbation on the microbial community composition and function in a tropical agricultural soil. *Environmental Science and Pollution Research*, 24:7139–7159.

Sghaier, H., Hezbri, K., Ghodhbane-Gtari, F., Pujic, P., Sen, A., Daffonchio, D., Boudabous, A., Tisa, L. S., Klenk, H. P., Armengaud, J., Normand, P., and Gtari, M. (2016). Stone-dwelling actinobacteria blastococcus saxobsidens, modestobacter marinus and geodermatophilus obscurus proteogenomes. *ISME Journal*, 10:21–29.

Shange, R. S., Ankumah, R. O., Ibekwe, A. M., Zabawa, R., and Dowd, S. E. (2012). Distinct soil bacterial communities revealed under a diversely managed agroecosystem. *PLoS ONE*, 7.

Sikorski, J., Baumgartner, V., Birkhofer, K., Boeddinghaus, R. S., Bunk, B., Fischer, M., Fösel, B. U., Friedrich, M. W., Göker, M., Hölzel, N., Huang, S., Huber, K. J., Kandeler, E., Klaus, V. H., Kleinebecker, T., Marhan, S., von Mering, C., Oelmann, Y., Prati, D., Regan, K. M., Richter-Heitmann, T., Rodrigues, J. F., Schmitt, B., Schöning, I., Schrumpf, M., Schurig, E., Solly, E. F., Wolters, V., and Overmann, J. (2022). The evolution of ecological diversity in acidobacteria. *Frontiers in Microbiology*, 13.

Song, W., Wang, Y., Peng, B., Yang, L., Gao, J., and Xiao, C. (2023). Structure and function of microbiomes in the rhizosphere and endosphere response to temperature and precipitation variation in inner mongolia steppes. *Frontiers in Plant Science*, 14.

Sur, S., Saha, S., Tisa, L. S., Bothra, A. K., and Sen, A. (2013). Characterization of pseudogenes in members of the order frankineae. *Journal of Biosciences*, 38:727–732.

Sáenz, J. S., Airo, A., Schulze-Makuch, D., Schloter, M., and Vestergaard, G. (2019). Functional traits co-occurring with mobile genetic elements in the microbiome of the atacama desert. *Diversity*, 11.

Wang, H., Bu, L., Tian, J., Hu, Y., Song, F., Chen, C., Zhang, Y., and Wei, G. (2021). Particular microbial clades rather than total microbial diversity best predict the vertical profile variation in soil multifunctionality in desert ecosystems. *Land Degradation and Development*, 32:2157–2168.

Waschulin, V., Borsetto, C., James, R., Newsham, K. K., Donadio, S., Corre, C., and Wellington, E. (2022). Biosynthetic potential of uncultured antarctic soil bacteria revealed through long-read metagenomic sequencing. *ISME Journal*, 16:101–111.
